# Tension sensing by FAK governs nuclear mechanotransduction, endothelial transcriptome and fate

**DOI:** 10.1101/2023.04.24.538195

**Authors:** Md Zahid Akhter, Pascal Yazbeck, Mohammad Tauseef, Mumtaz Anwar, Faruk Hossen, Sayanti Datta, Vigneshwaran Vellingiri, Jagdish Chandra Joshi, Nityanand Srivastava, Stephen Lenzini, Guangjin Zhou, James Lee, Mukesh K Jain, Jae-Won Shin, Dolly Mehta

**Author notes:** Lead contact: Dolly Mehta, Ph.D Professor & Interim Head Department of Pharmacology & Regenerative Medicine University of Illinois College of Medicine 835 S Wolcott Avenue Chicago, IL 60612.

## Abstract

Vascular endothelium forms a restrictive barrier to defend the underlying tissue against uncontrolled influx of circulating protein and immune cells. Mechanisms that mediate the transition from restrictive to leaky endothelium, a hallmark of tissue injury exemplified by acute lung injury (ALI), remain elusive. Using endothelial cell *(EC)-Fak^-/-^*mice, we show that FAK sensing and transmission of mechanical tension to the EC nucleus governs cell fate. In FAK- deleted EC, increased EC tension induced by Rho kinase caused tyrosine phosphorylation of nuclear envelope protein, emerin at Y74/Y95, and its localization in a nuclear cap. Activated emerin stimulated DNMT3a activity and methylation of the *KLF2* promoter, impairing the restrictive EC transcriptome, including *S1PR1*. Inhibiting emerin phosphorylation or DNMT3a activity enabled KLF2 transcription of *S1PR1*, rescuing the restrictive EC phenotype in *EC-Fak^-/-^* lungs. Thus, FAK sensing of tension transmission to the nucleus is crucial for maintaining a restrictive EC fate and lung homeostasis.

## Introduction

The vascular endothelium lining blood vessels plays a vital role in controlling the passage of plasma proteins and blood cells to the underlying tissue to achieve tissue-fluid homeostasis (Mehta and Malik, 2006). Endothelial cell (EC) attachment to the extracellular matrix (ECM) and adjacent cells generates a mechanical environment that maintains a restrictive barrier (Ingber, 2002; Mehta and Malik, 2006; Wang et al., 2009). A crucial but poorly understood aspect of these cells is their intrinsic property to oppose actin-myosin-induced intracellular tension (Daneshjou et al., 2015; Ingber, 2002), which may prevent their conversion from restrictive to leaky EC phenotype, a hallmark of tissue injury. Identifying these tension sensing mechanisms in EC is critical to reversing the leaky EC fate and inflammatory diseases such as acute lung injury (ALI) and acute respiratory distress syndrome (ARDS), and COVID-ARDS (Hariri and Hardin, 2020; Matthay and Zemans, 2011).

Focal adhesion kinase (FAK) is a ubiquitously expressed non-receptor tyrosine kinase that regulates several EC functions related to restrictive barrier formation. FAK links EC with the ECM by forming focal adhesions (Mehta and Malik, 2006; Parsons, 2003). FAK promotes EC proliferation and migration (Braren et al., 2006) and forms intercellular adhesions (Holinstat et al., 2006; Knezevic et al., 2009; Mehta and Malik, 2006; Quadri, 2012; Rajput et al., 2013). FAK also seems to serve as a mechanosensor downstream of fibronectin-linked integrin(Seong et al., 2013). Unlocking the autoinhibitory FAK-FERM domain using computational modeling and molecular dynamics of FAK showed that FAK directly senses the force perturbations (Zhou et al., 2015). We showed that conditional deletion of FAK in EC of adult mice (*EC-Fak^-/-^)* severely damaged blood vessels, causing ALI (Schmidt et al., 2013). LPS, a cell-wall component of gram-negative bacteria that causes ALI, also decreased FAK expression in adult lung vasculature (Schmidt et al., 2013; Ying et al., 2018).

We thus tested the hypothesis that FAK serves as an intrinsic tension sensing mechanism in EC to maintain EC restrictive fate. Using the *EC-Fak^-/-^* mouse model in combination with a multipronged approach, we identified FAK sensing of intracellular tension in EC as a critical determinant of nuclear mechanotransduction, which instructs the EC transcriptome in favor of a restrictive phenotype. We show that this function of FAK involves suppression of activities of the nuclear envelope protein, emerin, and DNMT3a to maintain KLF2 transcription of restrictive EC fate and lung vascular homeostasis.

## Results

### FAK softens EC to maintain KLF2 expression and lung vascular homeostasis

While the transition of EC from soft to stiff state disrupts endothelial barrier function, the mechanism remains unclear (Bastounis et al., 2019). We assessed if FAK was required to soften EC to maintain lung homeostasis. We used atomic force microscopy (AFM) to quantify intracellular tension, i.e., Young’s modulus (*E)* in control inactivated versus FAK depleted EC (Discher et al., 2005; Wong et al., 2020). We found that inactivated EC generates physiologically relevant intracellular EC tension ⁓8 KPa (**Fig. 1A and Supplementary Fig. 1A-B**). However, EC tension increased significantly in FAK-depleted EC (**Fig. 1A and Supplementary Fig. 1A-B**). Because FAK suppresses RhoA and myosin light chain phosphorylation (Holinstat et al., 2006; Mehta et al., 2002; Schmidt et al., 2013), in other studies, we determined the effect of overexpression of WT-RhoA, constitutively active (CA) RhoA (Q63L) or dominant-negative (DN) RhoA (T19N) mutants in regulating intracellular EC tension. EC expressing WT or CA-RhoA developed more tension than the DN-RhoA-EC **(Supplementary Fig. 1C)**. Consistently, inhibition of myosin-II activity (Even-Ram et al., 2007) or Rho kinase activity (Watanabe et al., 2007) in FAK depleted EC reversed the tension to the level seen in control EC **(Fig. 1B)**, indicating the requirement of FAK in maintaining the intracellular tension in EC by suppressing RhoA-Rho kinase pathway.

**Figure 1:**
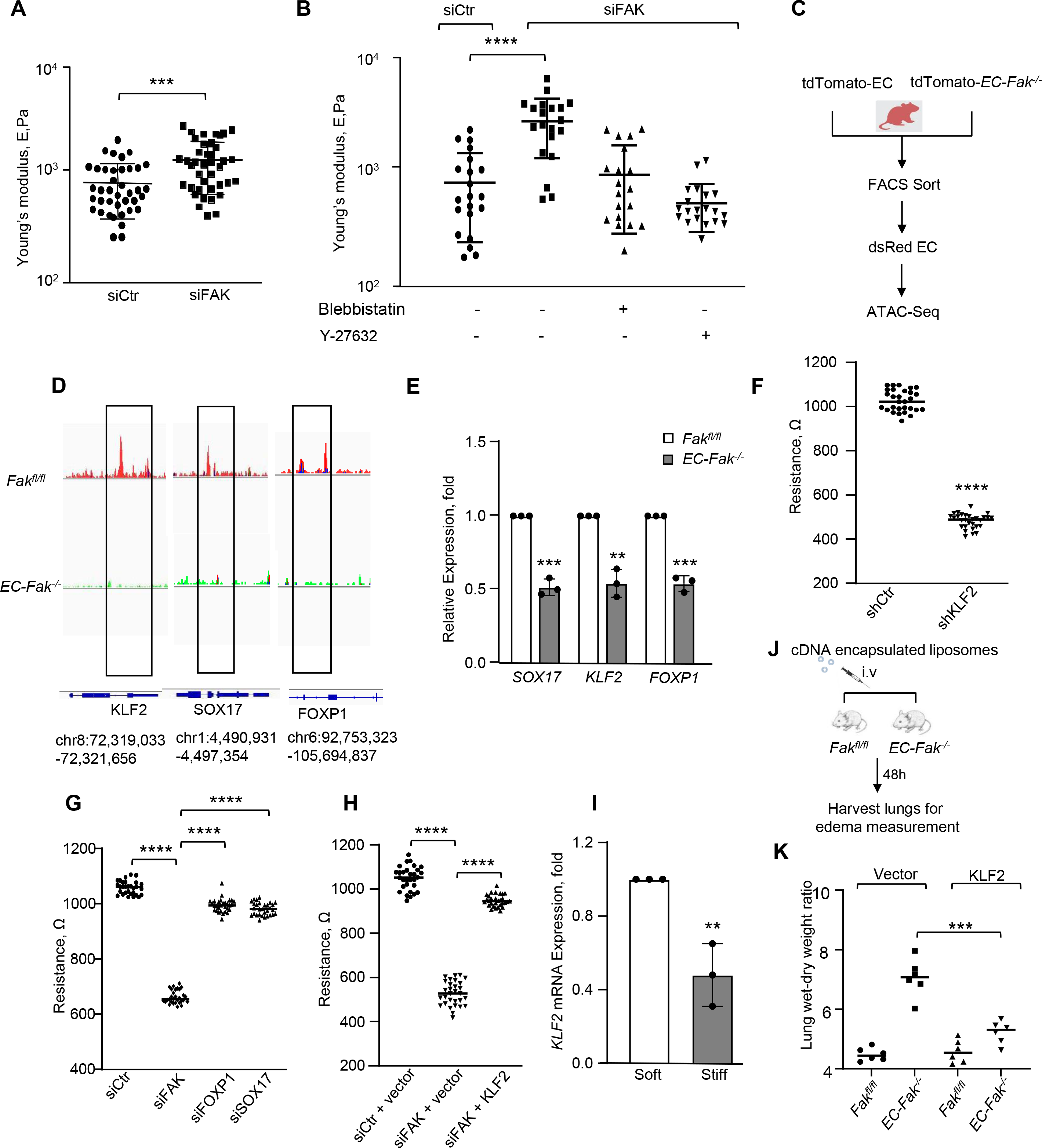
FAK regulates mechanotransduction and chromatin accessibility of transcription factors. A, Young’s modulus (E) was measured in a single control or FAK-depleted EC using atomic force microscopy (AFM) by nanoscale indentation at 1μ/s. N=40 cells/group, pooled from three independent experiments. **B,** FAK-depleted EC were treated with 20 µM of blebbistatin or 10 µM of Y- 27632 for 1 h. Young’s modulus (E) was then measured as described in **A**. The data were pooled from three independent experiments. **C,** Schematic diagram of EC sorting (CD31^+^CD45^-^) in lungs of tdTomato-*EC-FAK^-/-^*and control tdTomato mice for ATAC-seq analysis. **D,** Snapshot of genomic loci shows chromatin-accessible peaks at the transcription start sites (TSSs) for *KLF2*, *SOX17*, and *FOXP1* in FAK^+^ EC. Integrative Genomics Viewer (Broad Institute) was used to visualize the track peaks. **E,** mRNA expression of *SOX17, KLF2* and *FOXP1* in EC sorted from control or *EC-Fak* null mice lungs using qPCR. GAPDH was used as the house keeping gene (n=3). **F-G,** EC plated on TEER electrodes were transfected with either scrambled or *KLF2* shRNA or control and *SOX17*, *FOXP1*, and *FAK* siRNA. Basal barrier function was investigated 48 h after depletion of *KLF2*, *FAK*, *SOX17* and *FOXP1* (*n* = 8/group). The assay was repeated three times independently. **H,** TEER was measured as described in G in FAK-depleted EC after transducing vector, KLF2 cDNA post 48 h of FAK depletion (*n* = 8/group). Basal barrier function was assessed 72 h post-transfection. **I,** mRNA expression of *KLF2* in EC seeded on soft (⁓3 KPa) and stiff (⁓20 KPa) polyethylene glycol (PEG) hydrogels. GAPDH was used as internal control (n=3). **J,** Workflow of KLF2 cDNA delivery in EC of EC-FAK-null mice using liposomes. KLF2 cDNA was cloned in a VE-cadherin driven promoter to induce KLF2 expression specifically in EC. **K,** Lung edema was assessed by measuring wet-to-dry weight ratio of the lungs of indicated mice receiving either vector or KLF2 cDNA constructs using liposomes after 48 h (n=6 mice/group). Data in plots **A, B, E, F, G, H, I and K** show mean ± SEM. Statistical significance was assessed by One-way ANOVA followed by post hoc Tukey’s test for **B, G, H and K,** while unpaired t test was used for **A, E, F and I**. **** *P*< 0.0001, *** *P*< 0.001 and ** *P*< 0.01 relative to siCtr (A, B, F, G, H), *Fak^fl/^*^fl^ (E) *EC-Fak^-/-^* receiving vector (K). Also see Supplementary Fig. 1 and 2.

Mechanical tension influences proper chromatin assembly and gene transcription to regulate cell fate (Uhler and Shivashankar, 2017). To address the impact of FAK maintenance of EC tension in regulating chromatin accessibility to transcription factors and their roles in governing EC fate, we sorted EC from the lungs of control (tdTomato) and tamoxifen-inducible EC-specific FAK-null mice (tdTomato-*EC-Fak^-/-^*mice), generated as described (Akhter et al., 2021; Liu et al., 2019; Schmidt et al., 2013; Tran et al., 2016). We performed Assay of Transposase Accessible Chromatin (ATAC) sequencing to identify the tension-sensitive transcription factors regulating FAK-driven cell responses **(Fig. 1C)**. Findings showed that the chromatin region linked with the *KLF2*, *SOX17*, and *FOXP1* transcription factors were accessible in FAK*^+^***-**EC. However, these regions were inaccessible in FAK*^-^* EC **(Fig. 1D and Supplementary Fig. 1D-E)**. We validated these findings using sorted FAK^-^ or FAK^+^ EC **(Fig. 1E)**. Similarly, FAK-depleted human EC showed reduced expression of *KLF2*, *SOX17*, and *FOXP1* **(Supplementary Fig. 1F),** showing that FAK is required to promote the opening of their promoters and gene synthesis.

We next depleted *KLF2*, *FOXP1*, and *SOX17* in EC and determined barrier function by measuring transendothelial electrical resistance (TEER) (Akhter et al., 2021) across the monolayer to identify the transcription factors that phenocopy the findings in FAK-depleted EC (Holinstat et al., 2006; Mehta et al., 2002; Schmidt et al., 2013). We found that depletion of *KLF2* markedly reduced TEER, i.e., disrupted EC barrier function much like that seen in FAK-depleted EC **(Fig. 1F and Supplementary Fig. 2A)**. In contrast, depletion of *SOX17* or *FOXP1* had a modest effect on TEER **(Fig. 1G and Supplementary Fig. 2B-C)**. Restoration of KLF2 expression in FAK-depleted EC rescued EC barrier function **(Fig. 1H and Supplementary Fig. 2D)**. In line with these observations, the rescue of *SOX17* expression barely affected EC barrier dysfunction in FAK- depleted EC (**Supplementary Fig. 2E-F)**.

To assess if matrix stiffness similarly alters KLF2 expression, we prepared soft (⁓3 KPa) versus stiff (⁓20 KPa) polyethylene glycol (PEG) hydrogels as described(Lenzini et al., 2021). We found that EC plated on stiff PEG showed reduced *KLF2* expression compared with EC plated on the soft PEG **(Fig. 1I)**. We, therefore, assessed the causal role of KLF2 downstream of FAK in resolving vascular injury in *EC-Fak^-/-^* mice. We complexed KLF2-cDNA driven by the Cdh5 (VE-cadherin) promoter in liposomes (Tauseef et al., 2012) to express *KLF2* only in EC of *EC-Fak^-/-^* mice and quantified lung edema. As in EC, rescuing *KLF2* expression in EC of EC-FAK-null lungs resolved lung edema **(Fig. 1J-K)**. Immunohistochemical analysis of lungs receiving KLF2- cDNA confirmed KLF2 delivery to lung vessels **(Supplementary Fig. 2G)**. Thus, genome-wide chromatin accessibility of lung vessels identified KLF2 as the tension-regulated transcription factor downstream of FAK.

### FAK maintains *KLF2* synthesis by suppressing methylation of KLF2 promoter by DNMT3a

*KLF2* promoter contains CpG islands between -524 to -343 in the promoter region **(Fig 2A)** which are the target of DNA methyltransferases (DNMTs) (Kumar et al., 2013). We tested the possibility that FAK maintained *KLF2* expression by suppressing methylation of *KLF2* promoter. Bisulfite conversion of genomic DNA shows methylation states of individual cytosines on individual DNA molecules (Li and Tollefsbol, 2011). We found that 31% of the DNA underwent bisulfite conversion in control EC **(Fig. 2B-C).** However, 72% of the DNA converted into bisulfite in FAK-depleted EC, thus showing that loss of FAK leads to 3-fold induction of DNA methylation **(Fig. 2B-C)**. Methylation-specific PCR data similarly revealed a higher methylated: unmethylated ratio in FAK-depleted EC resulting in a 3.5-fold increase in DNA methylation **(Fig. 2D-E)**.

**Figure 2:**
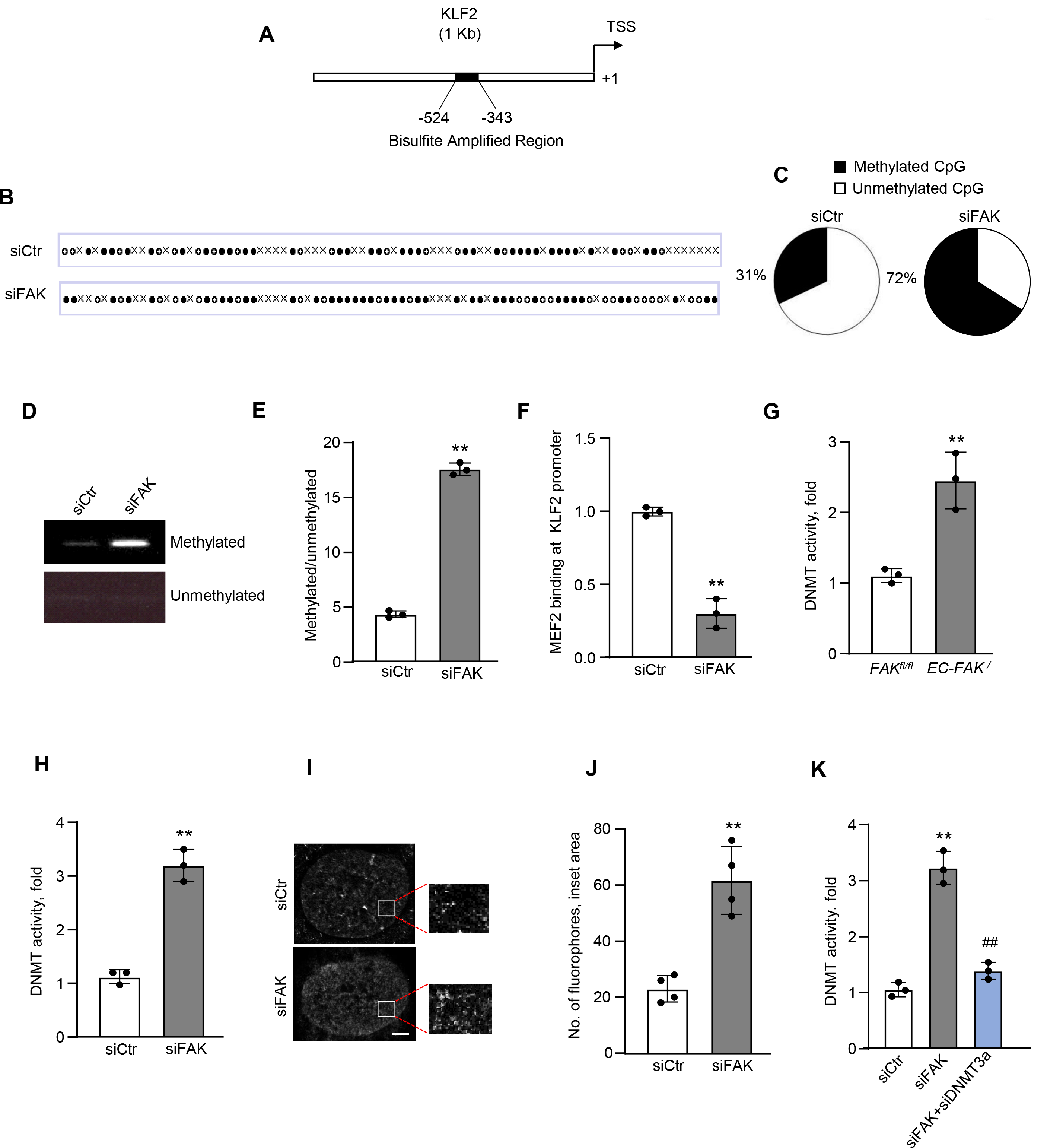
FAK upregulates KLF2 by suppressing DNA methylation of *KLF2* promoter. A, Schematic of *KLF2* promoter showing region (-343 to -524) containing CpG island. **B-C,** Bisulfite sequencing was performed on whole-genome DNA isolated from control or FAK-depleted EC following which subsequent cloning and sequencing was carried out. The methylation status of the *KLF2* promoter region was analyzed using QUMA (quantification tool for methylation analysis). Dark circle indicates methylated CpG and clear circle depicts unmethylated CpG. Pi-chart shows quantification of methylated and unmethylated CpG. **D-E,** Bisulfite conversion was performed on genomic DNA isolated from control or FAK-depleted EC. Methylation specific PCR (MS-PCR) was then carried out and the amplified product was quantified. **D** shows a representative DNA PAGE of amplified products while **E** shows densitometric analysis of the amplified products (n=3). **F,** Genomic DNA was isolated from control or FAK-depleted EC, and a chromatin immunoprecipitation (*ChIP*) assay followed by qPCR was performed to amplify MEF2 binding on *KLF2* promoter (*n* = 3). **G-H,** DNMT activity in *FAK^fl/fl^* and EC-FAK*-*null lungs **(G)** or control and FAK-depleted EC **(H)** (n=3). **I-J,** Representative **s**tochastic optical reconstruction microscopy (STORM) imaging of single nuclei in control versus FAK-depleted EC. EC were immunostained with anti-5mc antibody and AF-647 secondary antibody. Cells were imaged using GE OMX-SR Super-Resolution microscope (n=3). **I** shows representative diffraction corrected images. Indicated area is zoomed in inset. **J** shows quantitation of the 5-mc fluorophores. **K,** DNMT activity was measured in control EC, FAK-depleted EC, and EC depleted of both DNMT3a and FAK (n=3). Data plotted in **E, F, G, H, J and K** are given as mean ± SEM. Statistical significance was evaluated by one-way ANOVA followed by Post hoc Tukey’s test for K, while unpaired t-test was used for E, F, G, H and J. \*\**P*< 0.01 relative to siCtr (E, F, H, J, K) and *FAK^fl/fl^* (G). ## *P*< 0.01 relative to siFAK (K). Also see Supplementary Fig. 3.

MEF2 synthesizes *KLF2* by binding to the *KLF2* promoter at -733 bp (from TSS), the region containing CpG islands and sensitive to DNA methylation (Kumar et al., 2013). Chromatin immunoprecipitation (ChIP) analysis on the *KLF2* promoter showed a 70% decrease in MEF2 binding to the *KLF2* promoter in FAK-depleted EC as compared to control cells **(Fig. 2F and Supplementary Fig. 3A)**, corroborating the above findings that methylated *KLF2* promoter was inaccessible by its transcription factor impairing KLF2 synthesis.

DNMT1, DNMT3a, and DNMT3b process DNA methylation(Denis et al., 2011). DNMT1 is involved in hereditary DNA methylation, while DNMT3a and 3b regulate *de novo* DNA methylation (Denis et al., 2011). We found similar expressions of DNMT1, DNMT3a, and DNMT3b in *Fak^fl/fl^*or *EC-Fak^-/-^* lungs (**Supplementary Fig. 3B-C)**. However, DNMT3a activity increased by a factor of ∼2.5 in EC-Fak null lungs **(Fig. 2G)** and by a factor of ∼3-3.5 in FAK- depleted EC **(Fig. 2H)**. We also analyzed the nuclear distribution of 5-methylcytosine (5-mc), a measure of DNMT activity, at the single molecule level using specific 5-mc probes in conjunction with 3D stochastic optical reconstruction microscopy (STORM) in control versus FAK depleted EC. We found a ∼4-fold increase in 5-mc fluorophores in the nucleus of FAK-depleted EC than in control EC (**Fig. 2I-J**). We next knockdown DNMT3a or DNMT3b in FAK-depleted EC and found that depletion of DNMT3a but not DNMT3b (data not shown) rescued DNMT activity to near control levels **(Fig. 2K and supplementary Fig. 3D)**, suggesting that activated DNMT3a methylated *KLF2* promoter.

### Inhibition of DNMT3a activity reverses *KLF2* expression and EC fate from leaky to a restrictive phenotype

ALI is associated with decreased FAK and KLF2 expression (Schmidt et al., 2013; Ying et al., 2018). We therefore assessed if LPS increased DNMT activity in a WT mice and inhibition of DNMT3a activity would convert EC from leaky to restrictive phenotype via preserving KLF2 expression. We inhibited DNMT3a activity in mice specifically using theaflavin- 3,3’-digallate (TF-3) (1 mg/kg, *i.v*) (You et al., 2017) and after 1h induced endotoxemia (LPS, 10 mg/kg, *i.p*) (**Fig. 3A**). We found that LPS elevated DNMT activity **(Fig. 3B)**, which returned to the basal level after inhibiting DNMT3a activity **(Fig. 3B)**. As expected, LPS decreased *FAK* and *KLF2* expression in lungs **(Fig. 3C and supplementary Fig. 3E-F)**. However, LPS failed to decrease KLF2 expression in mice treated with TF3 and these mice rapidly resolved lung edema (**Fig. 3C-D**). Similarly, knockdown of DNMT3a in FAK depleted EC reversed *KLF2* expression to control levels **(Fig. 3E).** TF-3 and Pan-DNA methyltransferase inhibitor, 5-aza-2′- deoxycytidine (AZA) (Christman, 2002), rescued *KLF2* expression in FAK-depleted EC or EC- FAK-null lungs **(Supplementary Fig. 4A-C)**.

**Figure 3:**
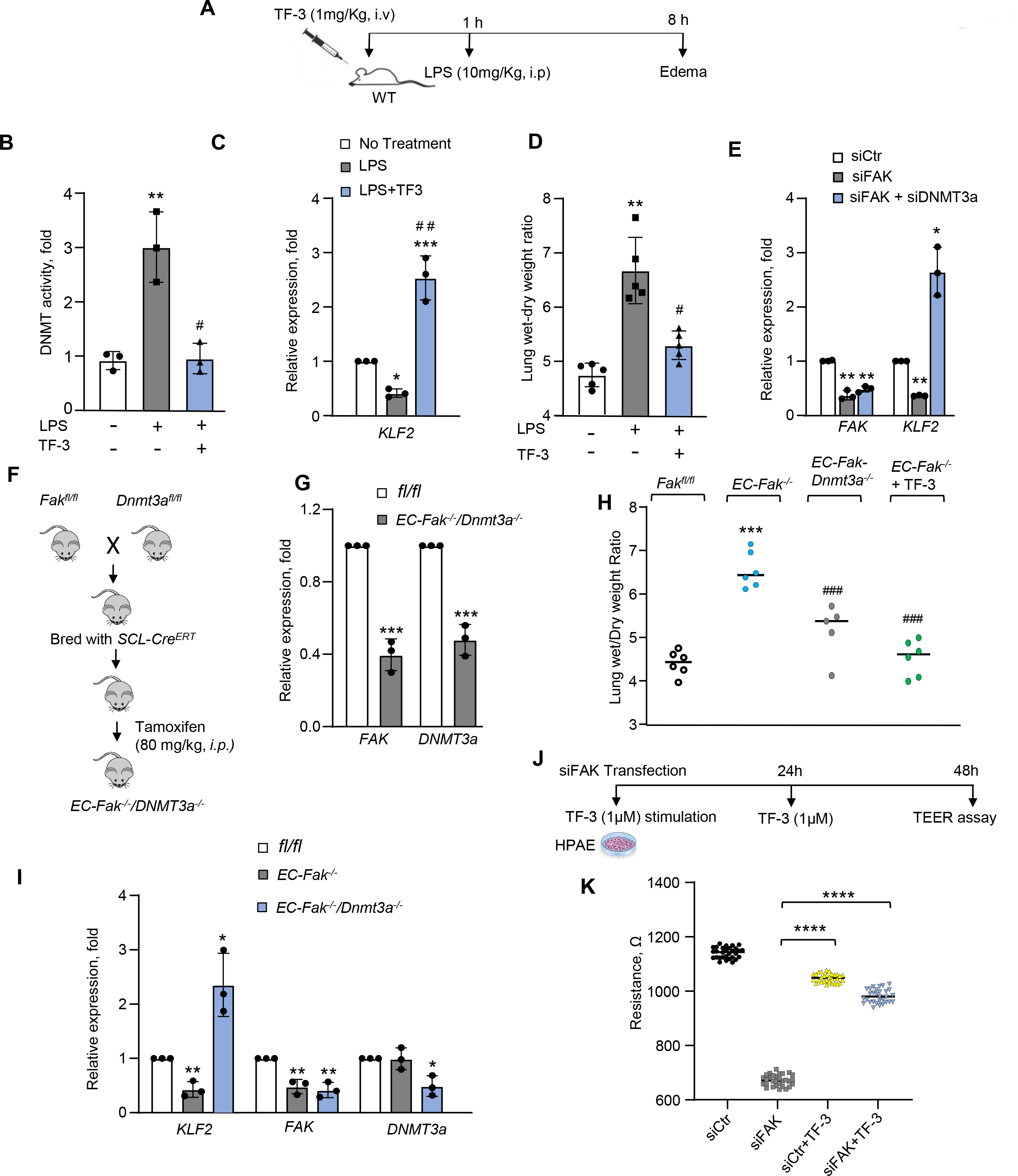
DNMT3a is required for LPS-induced lung injury. A, Protocol of inhibiting DNMT3a using TF-3 (1 mg/kg*, i.v*) in WT mice. After 1 h of TF-3 administration the mice received LPS (10 mg/kg, *i.p*) and lungs were harvested after 8 h (n=5mice/group). DNMT activity **(B)**, mRNA expression of *KLF2* **(C)** and lung edema **(D)** were determined (n=3). **E,** mRNA expression of indicated genes in control, FAK depleted, and FAK + DNMT3a-depleted EC. GAPDH was used as an internal control (n=3). **F,** Schematics showing generation of *EC-Fak/Dnmt3a^-/-^* (double knockout) mice. Offspring of *Fak^fl/fl^*and *Dnmt3a^fl/fl^* mice were bred with *SCL-Cre^ERT^*to generate *Fak/Dnmt3a^fl^SCL^CreERT^* mice. Tamoxifen (80 mg/kg, i.p) was injected into 4-week old *Fak/Dnmt3a^fl^SCL-^CreERT^*mice for five consecutive days followed by 7-day drug washout period. **G,** mRNA expression shows deletion of *FAK* and *S1PR1* in lungs from double knockout versus control mice. Experiments were repeated three times independently. **H,** Lung edema was assessed by measuring lung weight-dry weight ratio in *Fak^fl/fl^*, *EC-Fak^-/-^* and EC-FAK-null mice receiving TF-3 (1 mg/kg, i.v). **I,** mRNA expression of indicated genes in lungs of *Fak^fl/fl^*, *EC-Fak^-/-^* and *EC- Fak^-/-^/Dnmt3a^-/-^*. GAPDH was used as an internal control (n=3). **J-K,** TEER assay in FAK- depleted EC without or with TF-3 (1 µM) treatment as described in **J**. Data in plots **B, C, D, E, G, H, I and K** show mean ± SEM. Statistical significance was assessed by One-way ANOVA followed by post hoc Tukey’s test for **B, C, D, E, H, I and K,** while unpaired t test was used for **G**. \*\*\*\**P*< 0.0001 relative to siCtr (K), \*\*\**P*< 0.001 relative to no treatment (C), *Fak^fl/fl^* (G, H), \*\**P*< 0.01 to no treatment (B, D), siCtr (E) fl/fl (I), \**P*< 0.05 to siCtr (E), fl/fl (I). #*P*< 0.05 to LPS treated (B, D), ##*P*< 0.01 to no treatment (C) and ###*P*< 0.001 to *EC-Fak^-/-^*. Also see Supplementary Fig. 4.

If DNMT3a was responsible for subverting EC into leaky phenotype by downregulating *KLF2* and *S1PR1* expression, deletion of *DNMT3a* in EC of FAK-null mice should rescue gene expression and barrier function. We therefore generated inducible EC-specific double knockout (DKO) in which tamoxifen will delete both FAK and DNMT3a in *Fak/Dnmt3a^cre-scl-ERT^* (**Fig. 3F-G and supplementary Fig. 3D)**. In contrast to FAK, DNMT3a deletion alone in EC did not alter vascular permeability at homeostasis (data not shown). However, deletion of DNMT3a in EC of *EC-FAK^-/-^* restored the lung vascular barrier **(Fig. 3H)**. We also administered AZA (0.5 mg/kg, *i.v.*) and TF-3 (1 mg/kg, *i.v.*) in *EC-Fak* null mice, and after 24 h, we assessed vascular permeability. We found that inhibition of DNMT3a activity rescued vascular homeostasis in *EC-Fak^-/-^* mice **(Fig. 3H and supplementary Fig. 4E)**. TF-3 addition also rescued the endothelial barrier **(Fig. 3J-K)**.

To assess if EC stiffening also increased DNMT3a activity, we quantified DNMT activity in EC plated on stiff versus soft matrices. DNMT activity increased by 4-fold in EC plated on the stiff-matrix than in EC plated on the soft matrix **(Supplementary Fig. 4F)**. These findings, along with data in Fig. 1-2, demonstrate that EC stiffening is a generalized phenomenon during vascular injury that in turn upregulates DNMT3a activity to compromise KLF2 synthesis and thereby EC fate.

### KLF2 maintains restrictive EC fate by synthesizing S1PR1

We next sorted EC from control and EC-FAK-null lungs to assess the EC transcriptome dictating EC restrictive fate and whether it required KLF2 for gene synthesis. RNA-seq analysis in FAK^-^ EC identified *S1PR1*, a G-protein-coupled receptor that we previously showed maintains EC barrier function (Akhter et al., 2021), as the crucial gene reduced in these EC **(Fig. 4A)**. To test the hypothesis that KLF2 synthesis of *S1PR1* maintained EC restrictive phenotype downstream of FAK, we used multi-pronged approaches. First, we determined *S1PR1* expression at mRNA and protein levels in FAK-null EC and found these significantly lower than the control EC **(Fig. 4B-D)**. FAK-depleted human EC, similarly, showed significantly reduced expression of S1PR1 **(Supplementary Fig. 5A-B)**. However, FAK depletion did not affect other S1PR receptors known to occur in EC, such as S1PR2 and S1PR3 (**Supplementary Fig. 5C**). The expression of either S1P synthesizing enzymes, SPHK1 or SPHK2 or KLF4, which also maintains EC integrity, was not altered **(Supplementary Fig. 5C)**.

**Figure 4:**
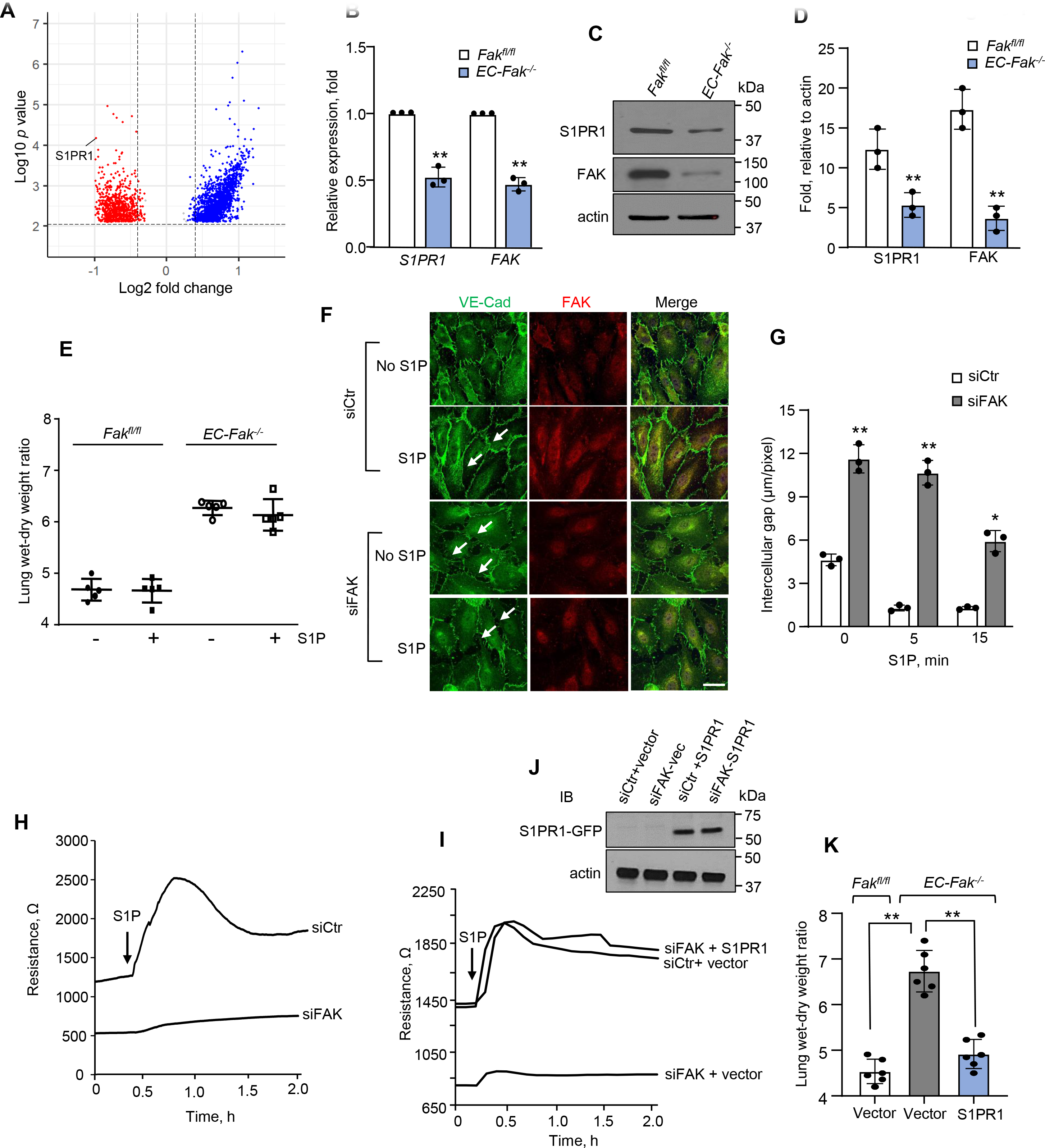
Transcriptome analysis identifies *S1PR1* as the gene of interest maintaining vascular integrity downstream of FAK. A, Volcano plot of differential genes from bulk RNA- seq analysis of EC sorted from *Fak^fl/fl^* and *EC-Fak^-/-^* mice lungs. *S1PR1* mRNA **(B)** or protein **(C)** expression in FAK^+^ versus FAK^-^ EC. GAPDH was used an internal control for mRNA while actin was used as a loading control (n=3). **D,** Corresponding densitometry of the blots. **E,** S1P (1 mg/kg, *i.v*) was administered to *Fak^fl/fl^* or *EC-Fak^-/-^*mice and after 30 min lung edema was assessed by measuring wet-to-dry weight ratio of the lungs (n=5 mice/group). **F,** A representative micrograph shows effect of S1P in re-assembling adherens junctions in control versus FAK-depleted EC. EC received 1 µM S1P for 15 min after which the cells were co-immunostained with anti-VE-cadherin and anti-FAK antibodies. Images were acquired using confocal microscope. *Note,* Control EC were plated at sub-confluent level to match gaps in FAK depleted EC before S1P stimulation (n=3). Scale bar 20 µm. **G,** Plot shows intercellular gap in control and FAK-depleted EC after with or without 1 µM S1P for 15 min. Gap area was assessed by ImageJ. The experiment was independently repeated three times. **H,** TEER in control and FAK-depleted EC at baseline and after addition of 1 µM S1P (*n* = 8 wells/group). The assay was repeated three times independently. **I-J,** EC plated on TEER electrodes were processed as in H for 48 h. EC were then re-transfected with vector or GFP-tagged WT-S1PR1 cDNA and TEER was measured **(I)** and S1PR1 expression **(J)** was quantified after 72 h after addition of 1 µM S1P (*n* = 8wells/group). The assay was repeated three times independently. Immunoblot of EC shows S1PR1 expression taking actin as a loading control (n=3). **K,** Edema measurement was performed in *Fak^fl/fl^* and *EC-Fak^-/-^* mice 48 h after delivery of liposomes containing VE-cadherin promoter driven GFP-tagged WT-S1PR1 cDNA as in 1K (n=6 mice/group). Data in plots **B**, **D**, **E**, **G and K** show mean ± SEM. Statistical significance was assessed by one-way ANOVA followed by Post hoc Tukey’s test in E and K, while unpaired t-test was used in B, D and G. ** *P*< 0.01 relative to *FAK^fl/fl^*(B, D) and *Fak^fl/fl^* and *EC-Fak^-/-^* receiving empty vector (K) * *P*< 0.05 relative to siCtr (G). Also see Supplementary Fig. 5.

S1P enhances basal barrier function in EC and repairs lungs from vascular injury (Akhter et al., 2021; Garcia et al., 2001; Tauseef et al., 2008). Thus, we determined whether S1P addition would promote endothelial integrity in FAK-null EC despite of ∼50% reduction in *S1PR1* expression. S1P failed to resolve vascular barrier leak in *EC-Fak^-/-^* mice **(Fig. 4E)**. We also determined the effect of S1P in strengthening endothelial monolayer integrity by quantifying inter-endothelial gap area and TEER in control monolayers and FAK-depleted EC. We found that S1P rapidly increased VE-cadherin accumulation at cell-cell junctions in control EC, thus decreasing the amount of intercellular gap area **(Fig. 4F-G)**. However, S1P failed to increase VE-cadherin accumulation in FAK-depleted EC, and as a result, the monolayer remained leaky **(Fig. 4F-G)**. As expected, S1P enhanced barrier function, i.e., TEER in control EC but not observed in FAK depleted EC **(Fig. 4H and Supplementary Fig. 5D)**.

Next, we restored *S1PR1* expression by transducing S1PR1 cDNA in FAK-depleted EC and found that restoration of *S1PR1* rescued basal and S1P-mediated increase in TEER **(Fig. 4I-J and Supplementary Fig. 5D)**. Based on these findings, we transduced S1PR1 cDNA driven by the Cdh5 promoter in EC of EC-FAK-null mice as above and found that transduction of S1PR1 in lungs reversed EC restrictive fate in FAK-null mice **(Fig. 4K and Supplementary Fig. 5E-F)**.

Because *in silico* analysis of the *S1PR1* promoter showed three KLF2 binding sites within the 1 kb promoter region of the mouse *S1PR1* gene(Skon et al., 2013) **(Fig. 5A)**, we co-transfected EC with the WT-S1PR1 luciferase promoter or mutated S1PR1 luciferase promoter along with KLF2 cDNA and determined promoter activity. KLF2 induced the *S1PR1* promoter activity in EC transducing the WT-S1PR1 promoter (**Fig. 5B**). However, KLF2 failed to increase the promoter activity in EC transducing the mutated S1PR1 promoter **(Fig. 5B)**. Further, depletion of KLF2 reduced *S1PR1* mRNA and protein expression **(Fig. 5C-E),** and S1P failed to enhance EC barrier function in KLF2-depleted EC **(Fig. 5F and supplementary Fig. 5G)**. Restoration of *KLF2* expression in FAK-depleted EC rescued *S1PR1* expression and S1P enhancement of barrier function **(Fig 5G-I)**. Also, inhibition of DNMT3a in FAK depleted EC reversed *S1PR1* expression **(Fig. 5J-L)**. These data show that KLF2 functioned downstream of FAK by synthesizing *S1PR1*.

**Figure 5:**
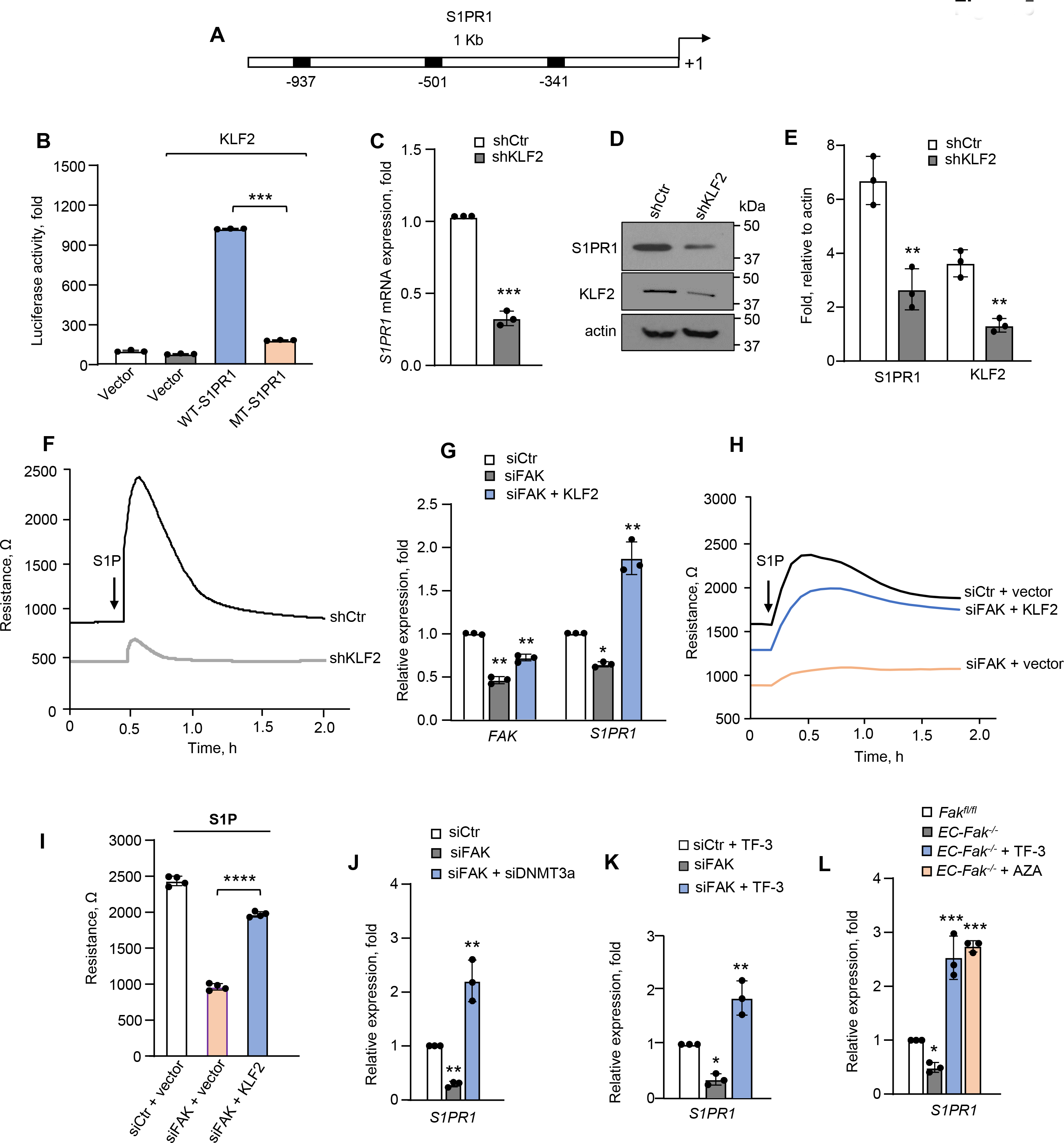
KLF2 regulates S1PR1 expression downstream of FAK. A, Schematic representation of *S1PR1* promoter with three KLF2 binding sites (-341, -501, and -937 from TSS). **B,** EC were co-transfected with WT-S1PR1 luciferase promoter or mutated S1PR1 luciferase promoter (which lacks three KLF2 binding sites) along with KLF2 cDNA (n=3). Luciferase activity was determined after 24 h. **C-D,** EC were transfected with control shRNA or KLF2 shRNA. After 48 h, *S1PR1* mRNA **(C)** or protein expression **(D)** was determined. GAPDH was used as a loading control for mRNA and actin was used as loading control for protein (n=3). **E,** Corresponding densitometry of the above blots. **F,** EC plated on TEER electrodes were transfected with control or KLF2 shRNA. After 48 h, cells were stimulated with S1P (1 µM) and TEER was measured as in 2G (n = 8wells/group). **G,** After 48 h of siFAK transfection, EC were co-transfected with vector or KLF2 cDNA. mRNA expression of indicated genes was assessed by qPCR. GAPDH was used an internal control (n=3). **H-I,** EC plated on TEER electrodes were transfected with control or FAK siRNA. After 48 h, cells were co-transfected with WT-KLF2 cDNA. After 72 h cells were stimulated with S1P, and TEER was assessed as in Fig. 1F (*n* = 8wells/group). **J-K,** *S1PR1* mRNA expression in FAK-depleted EC after DNMT3a knockdown **(J)** or inhibition with 1 µM theaflavin-3,3’-digallate (TF-3) for 4 h. **(K). L**, *S1PR1* mRNA in *EC-Fak^-/-^* mice after 24 h injection of AZA (0.5 mg/kg, *i.v.*) or TF-3 (1mg/kg, *i.v*). GAPDH was used an internal control (n=3). Data plotted in **B, C, E, G, I, J, K and L** are given as mean ± SEM. Statistical significance was assessed by one-way ANOVA followed by Post hoc Tukey’ test in B, G, I, J, K and L and unpaired t test was used in C and E. \*\*\*\**P*< 0.0001 relative to siFAK+KLF2 (I), \*\*\**P*< 0.001 to WT-S1PR1 (B), shCtr (C) *Fak^fl/fl^* (L), \*\**P*< 0.01 relative to shCtr (E), siCtr (G, J) siCtr+TF-3 (K) and \**P*< 0.05 siCtr (G), siCtr+TF-3 (K). Also see Supplementary Fig. 5-6.

### Emerin activates DNMT3a to induce a leaky EC phenotype

LINC (linker of nucleoskeleton and cytoskeleton) complex plays a critical role in sensing tension at the nuclear envelope (Tifft et al., 2009). Because emerin activation is linked with increase in nuclear stiffening (Guilluy et al., 2014), we tested the hypothesis that FAK maintains restrictive EC by suppressing emerin activity. We first immunostained control and FAK-depleted EC with anti-emerin antibody, which showed diffused emerin localization in control EC **(Fig. 6A-B).** However, in FAK-depleted EC, emerin condensed to form a nuclear cap **(Fig. 6A-B).** Similarly, emerin localized as a nuclear cap in EC plated on stiff matrix than a soft matrix **(Supplementary Fig. 6A-B)**.

**Figure 6:**
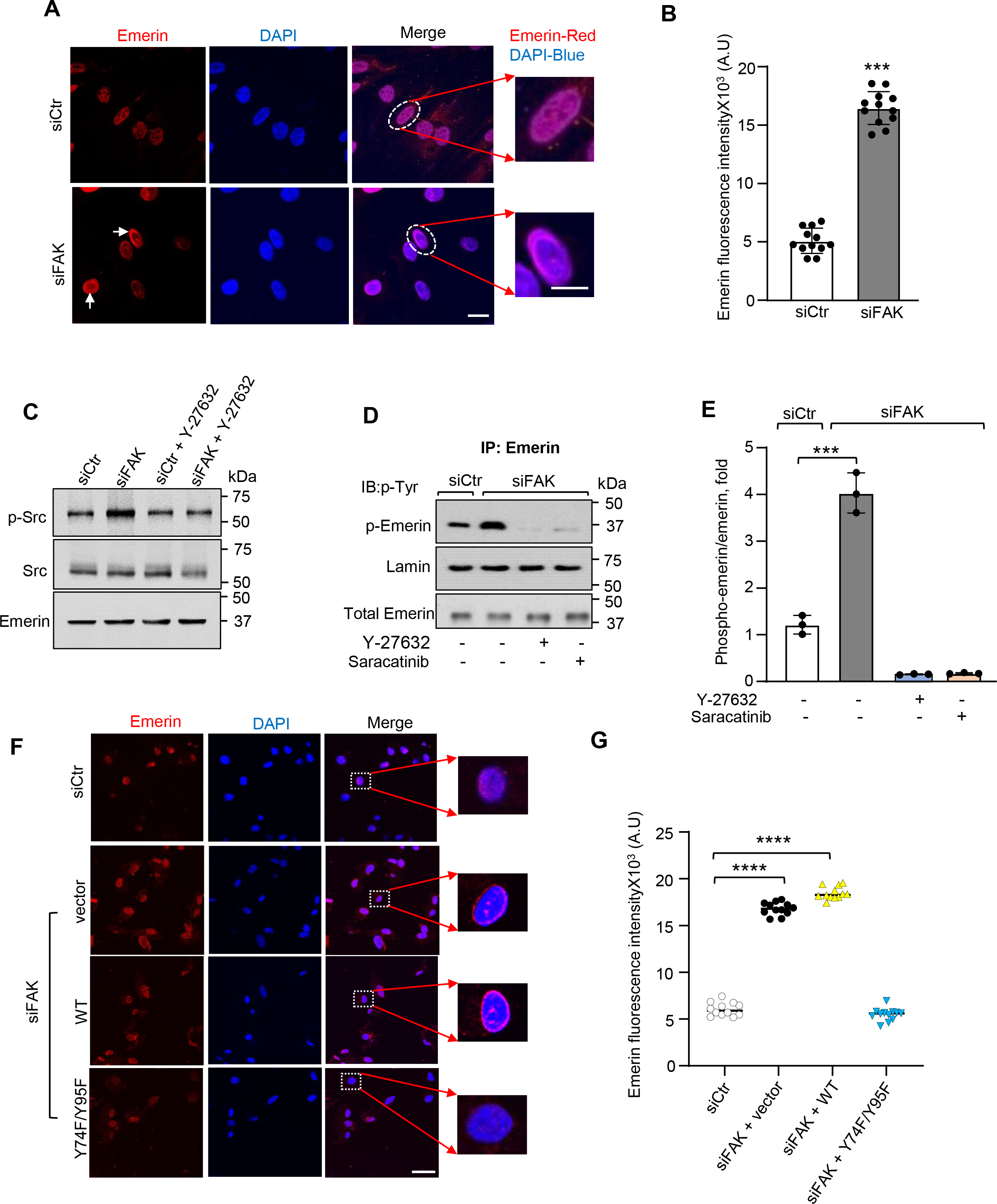
FAK regulates nuclear mechanotransduction via emerin: A, Representative micrograph shows emerin localization in nuclei in control versus FAK-depleted EC. The EC were stained with anti-emerin antibody and an appropriate fluorescent secondary antibody. Nuclei were labelled using DAPI to assess emerin localization. Boxed regions of emerin localization are magnified x3 on the right. Scale bar: 20 µm for unmagnified images and 60 µm for magnified images. **B,** Quantitation of emerin peripheral reorganization. An area of uniform size was drawn around the nuclear periphery in control or FAK-depleted EC and fluorescence intensity was quantified. Data is representative of at least n=12 from experiments that were repeated three times independently. **C,** Control or FAK-depleted EC were treated with a ROCK inhibitor as in Fig 1B and nuclei were isolated and immunoblotted with anti-phospho-cSrc and anti-pan-cSrc antibodies. Immunoblot with anti-emerin antibody was used to confirm the nuclear fraction. A representative blot is shown from experiments that were repeated three times independently. **D-E,** Nuclear fraction obtained from control or FAK-depleted EC were treated with Y-27632 (10 μM) as in **A** or cSrc inhibitor (Saracatinib, 100 nM) for 2 h were immunoprecipitated with an anti-emerin antibody. Immunocomplexes were immunoblotted with p-Tyr (PY99/PY29, 1:250 dilution) to assess emerin tyrosine phosphorylation over total emerin. **D,** shows a representative blot while **E** shows densitometry from experiments that were repeated three times. **F-G,** Micrograph shows changes in emerin peripheral reorganization in FAK-depleted EC transducing WT-or a phosphodefective emerin mutant (Y74F/Y95F) as described in **A.** Scale bar 20 µm. Fluorescence intensity of emerin was assessed as in **B**. Data plotted in **B, E** and **G** are given as mean ± SEM. Statistical significance was assessed by one-way ANOVA followed by Post hoc Tukey’s test in E and G and unpaired t-test was used in B. \*\*\*\**P*< 0.0001, \*\*\**P*< 0.001 relative to siCtr (G, E, B). Also see Supplementary Fig. 6.

Emerin undergoes tyrosine phosphorylation via tyrosine kinases such as cSrc upon sensing tension (Guilluy et al., 2014; Tifft et al., 2009). cSrc activation compensates for loss of FAK function (Sieg et al., 1998). Thus, we next assessed if loss of FAK in EC activated emerin by inducing post-translational modification via cSrc. We found that FAK depletion increased cSrc phosphorylation at Y419, a measure of cSrc activity (Higuchi et al., 2021)**(Fig. 6C and Supplementary Fig. 6C)**. Rho kinase inhibition reversed cSrc activity as in control cells (**Fig. 6C and Supplementary Fig. 6C**). Compared to control EC, FAK depletion, therefore, enhanced emerin tyrosine phosphorylation by ∼4 fold **(Fig. 6D-E)**. Inhibition of Rho kinase and cSrc restored emerin phosphorylation in FAK-depleted cells equivalent to control cells **(Fig. 6D- E)**. Together, the above findings identified emerin as a novel target of FAK and suggested that FAK control of intracellular tension transmission to the nuclear envelope is required to suppress emerin activation.

Emerin tyrosine phosphorylation at Tyr 74 and Tyr 95 regulates nuclear stiffening (Guilluy et al., 2014). Therefore, we examined the effects of modulating emerin activity in FAK-depleted cells by disrupting emerin phosphorylation at these residues (Y74F/Y95F) on DNMT activity, gene expression, and EC fate. First, we overexpressed mutated (Y74F/Y95F) or WT emerin in FAK depleted EC and assessed emerin organization. We found that mutated but not the WT- emerin reversed emerin expression in FAK-depleted EC from nuclear cap to diffuse one **(Fig. 6F- G)**. We next investigated whether enhanced phosphorylation of emerin at Tyr 74 and 95 increased DNMT3a activity in FAK-depleted EC. We found that Inhibition of emerin phosphorylation rescued DNMT activity in FAK-depleted cells as in control cells **(Fig. 7A and Supplementary Fig. 6D)**. This response was not seen in FAK-depleted EC expressing WT-emerin **(Fig. 7A and Supplementary Fig. 6D)**. Transduction of phosphodefective-emerin (Y74F/Y95F) restored *KLF2* or *S1PR1* expression and reduced intercellular gaps in FAK-depleted EC **(Fig. 7B- D)**. However, WT emerin failed to reverse these responses in FAK-depleted EC **(Fig. 7B-D)**. In line with these findings, emerin mutant restored the monolayer resistance in FAK depleted EC **(Fig. 7E)**. The results suggest that in the absence of FAK, phosphorylated emerin plays a critical role in mediating DNMT3a activity that suppresses *KLF2* synthesis of *S1PR1* leading to leaky EC fate.

**Figure 7:**
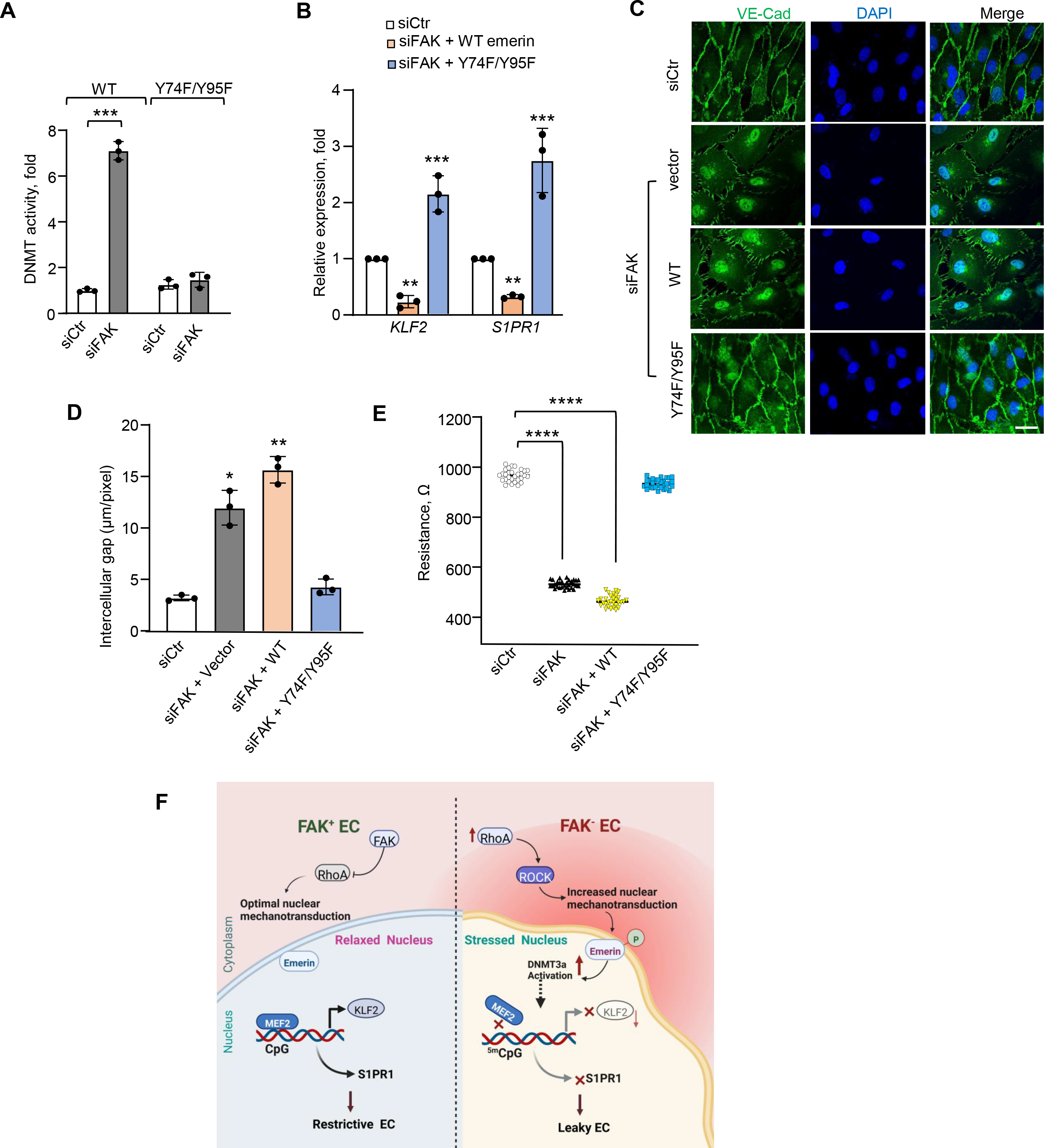
Suppression of emerin activity converts leaky to restrictive EC. A, DNMT activity was measured in FAK depleted EC transduced with WT-emerin or phosphodefective emerin mutant after 24 h post-transfection as in Fig. 2G (n=3). **B,** mRNA expression of *KLF2* and *S1PR1* in FAK depleted EC transducing WT-or phosphodefective-emerin. GAPDH was used as an internal control (n=3). **C,** Representative micrograph demonstrates intercellular gap as measured by immunostaining with anti-VE-Cadherin antibody in FAK-depleted EC after transduction with WT- or phosphodefective-emerin mutant. Scale bar 20 µm. **D,** Intercellular gap was measured as in **4F. E,** EC plated on TEER electrodes were transfected with either Ctr or FAK siRNA and after 48h WT- or phosphodefective- emerin mutant was transduced. TEER was assessed after 72 h (*n* = 8/group). The assay was repeated three times independently. Data plotted in A, B, D and E are given as mean ± SEM. Statistical significance was assessed by one-way ANOVA followed by Post hoc Tukey’s test. \*\*\*\**P*< 0.0001 relative to siCtr (E), \*\*\**P*< 0.001 siCtr (A, B), \*\**P*< 0.01 siCtr (D), \**P*< 0.05 siCtr (D). **F,** Model of FAK suppression of nuclear mechanotransduction and synthesis of genes maintaining vascular barrier. We posit that loss of FAK increases RhoA activity which in turn increases intracellular mechanical tension by activating Rho kinase and myosin ATPase activity. Increased tension activates cSrc which phosphorylates emerin. Phosphorylated emerin activates DNMT3a, which in turn methylates the *KLF2* promoter, blocking binding of its transcription factor, MEF2, and hence KLF2 synthesis. Suppression of KLF2 synthesis compromises *S1PR1* transcription leading to conversion of restrictive EC to leaky EC which impairs vascular homeostasis.

## Discussion

The mechanisms controlling the re-establishment of the EC restrictive fate remain the key to reversing diseases caused by leaky vasculature syndrome, including acute lung injury (Bhattacharya and Matthay, 2013; Mehta and Malik, 2006). In this study, we have demonstrated FAK as a tension sensor and guardian of the EC transcriptome and phenotype. We showed that loss of FAK in EC increased intracellular tension impinging on the nucleus leading to emerin reorganization and tyrosine phosphorylation at Y74/Y95 in a Rho-kinase-cSrc dependent manner. Phosphorylated emerin upregulated DNMT3a activity leading to the methylation of the *KLF2* promoter and suppression of *S1PR1* synthesis, and conversion of EC to their leaky phenotype. Therefore, impairment of emerin and DNMT3a function rescued the synthesis of *KLF2* and its transcriptional activity in EC of *EC-Fak^-/-^* lungs and restored lung homeostasis. In the WT endotoxemia mouse model, impairing DNMT3a activity also resolved lung injury by upregulating *KLF2*. While previous studies demonstrated FAK to be a sensor of mechanical tension (Li et al., 2002), our present findings point to the impact of FAK mechanosensing and transmission of the tension to the nucleus on the activation of *KLF2* transcription and acquisition of restrictive EC fate for lung fluid balance.

The mechanical environment critically regulates cell fate (Ingber, 2002; Wang et al., 2009). Actin stress fiber formation, a measure of EC contraction, increased upon elevation of substrate stiffness from a physiologically relevant level (8.6 kPa) to a non-physiological level (42 kPa) (Birukova et al., 2013). Similarly, oscillatory but not laminar flow, increased EC permeability(Sei et al., 2017). However, the mechanism and the magnitude of the intracellular tension in a restrictive versus a leaky phenotype remain elusive. We showed that Young’s modulus in control EC was around ∼800 Pa, which increased significantly in FAK-depleted EC. Pulmonary EC shows a wide variation in Young’s modulus (250-300 Pascal to 0.1 Megapascal) (Birukova et al., 2009). The values we obtained in the current study using AFM at an indentation rate of 1 m/s fall near the lower limit of the previously reported range (Delgadillo et al., 2020). FAK regulates focal adhesion turnover and orchestrates outside-in signaling (Braren et al., 2006; Li et al., 2002; Mehta and Malik, 2006; Parsons, 2003). Thus, FAK deletion may increase intracellular tension passively. We showed that inhibiting Rho kinase and myosin ATPase activity was sufficient to rescue EC tension to normal levels in FAK-depleted EC. Therefore, we have demonstrated that loss of FAK increases EC tension due to activation of Rho-kinase ATPase downstream of Rho-kinase, which we proved to be an essential factor in the genesis of the observed phenotypes in FAK-depleted EC.

Matrix rigidity is known to regulate the transmission of intracellular tension to the nucleus(Hoffman et al., 2011). While the details of the signaling pathway remained elusive, cells plated on a stiff matrix, but not a soft matrix, showed nuclear stiffening (Guilluy and Burridge, 2015). An externally applied mechanical load or fibronectin sensed by integrins at the focal adhesions activates FAK, cSrc, and RhoA signaling (Daneshjou et al., 2015; Li et al., 2002; Tomakidi et al., 2014). We showed that depletion of FAK in EC augmented intracellular force, cSrc phosphorylation, DNMT3a activity, and emerin activation, which also phenocopied in stiff EC. We, therefore, conclude that FAK controlled tension development and tension transmission to the nucleus by maintaining matrix rigidity and RhoA-cSrc activities.

Several transcription factors such as ERG (Lathen et al., 2014), FoxC (Xia et al., 2021), KLF2 (Atkins and Jain, 2007), Sox17 (Liu et al., 2019), and EGR1 (Akhter et al., 2021) are known to be involved in maintaining the barrier restrictive EC transcriptome. However, our findings identified KLF2 as the tension-regulated transcription factor using global ATAC-seq. EC express KLF2 and KLF4 at high levels, which regulate genes crucial for EC adaptation to sheer flow and barrier function (Atkins and Jain, 2007). We showed that FAK deletion markedly suppresses *KLF2* activity but had no effect on *KLF4* expression, indicating KLF2 to be the essential kruppel-like transcription factor dysregulated upon FAK deletion leading to impaired synthesis of the EC transcriptome. In line with this notion, rescuing KLF2 expression in FAK-depleted EC or lung vessels restored the restrictive EC phenotype and, thereby, lung homeostasis.

Specific open-chromatin states in regions recognized by epigenetic regulators, including DNA methyltransferases, enable the synthesis of transcription factors driving EC fate during development (Russell-Hallinan et al., 2021). Moreover, DNA methylation regulates KLF2 expression (Kumar et al., 2013). Using ATAC-seq analysis and methylation profiling, we discovered that FAK-depleted EC has a leaky phenotype due to methylation of the *KLF2* promoter. The methylated *KLF2* was not accessed by its transcription factor, MEF2, leading to reduced *KLF2* synthesis. Our data showed that methylation of the *KLF2* promoter and downregulation of KLF2 was due to activation of DNMT3a. Hence, deletion of DNMT3a in EC of FAK-null mice or its inhibition rescued *KLF2* and *S1PR1* synthesis and hence lung vascular homeostasis.

KLF2 regulates several EC genes (SenBanerjee et al., 2004), but our transcriptome analysis of FAK-null EC identified *S1PR1* as the top hit. *S1PR1* maintains vascular homeostasis basally and reverses vascular barrier function after lung injury(Akhter et al., 2021; McVerry et al., 2004). We showed that loss of FAK suppresses S1PR1 protein and mRNA expression due to impaired KLF2 activity. Furthermore, we showed that KLF2 bound the *S1PR1* promoter and increased S1PR1 luciferase activity. Thus, transduction of KLF2 in EC of FAK-null mice restored lung vascular homeostasis. In line with these findings, KLF2 induced *S1PR1* expression in response to statins in lung EC and T cells(Skon et al., 2013), leading us to propose that downstream of FAK, transcriptional regulation of *S1PR1* by KLF2 is critical to the establishment of the lung EC fate in favor of the restrictive phenotype.

Phosphorylation of emerin’s Y74 and Y95 residues by cSrc mediates mechanical adaptation of nuclei to mechanical force(Guilluy et al., 2014). Previously, we showed that FAK depletion induces RhoA activity leading to disruption of EC barrier function(Holinstat et al., 2006; Schmidt et al., 2013). In the present study, we showed that Rho kinase activation of myosin-ATPase activity caused a rise in intracellular tension and emerin activation, converting EC from the restrictive to leaky phenotype. Overexpression of a phosphodefective emerin in FAK-depleted EC suppressed DNMT activity and reversed KLF2 and S1PR1 expression in FAK-depleted EC.

The mechanism by which emerin activates DNMT3a needs parsing parsed out. A possible scenario may be that emerin-mediated nuclear stiffening alters histones leading to activation of DNMT3a activity (Guo et al., 2015). Nevertheless, our findings support recent studies, which suggest that DNMT3a plays a crucial role in suppressing KLF2 expression in EC under flow conditions (Parmar et al., 2006). These findings indicate that transmission of increased intracellular force to the nucleus following barrier disruption induces the EC phenotype switch. Indeed, we showed that overexpression of phosphodefective emerin rescued the synthesis of the restrictive EC transcriptome and stabilized barrier function in FAK-depleted EC, thus demonstrating emerin to be the target of FAK and its impact on EC fate and vascular homeostasis.

Together our evidence provides crucial insights into how FAK shapes the EC transcriptome and prevents its conversion to the injury-inducing EC phenotype. These findings establish suppression of KLF2 transcriptional activity to be the target of aberrant nuclear mechanotransduction and the leaky EC phenotype in the absence of FAK **(Fig. 7F)**. Acute lung injury and ARDS remain challenging lung diseases despite the emergence of new pharmacological and cell therapies (Matthay and Zemans, 2011). An important question is whether EC can be converted from their leaky to restrictive phenotype during injury to promote vascular repair and restore lung homeostasis. We showed that inhibiting emerin phosphorylation or DNMT3a activity enabled KLF2 transcription of *S1PR1* and rescued the EC-restrictive phenotype. These findings suggest that FAK is a potential target for restoring the EC restrictive phenotype, vascular homeostasis, and tissue function.

### STAR methods

Sphingosine-1-phosphate (BML-SL140-001) was purchased from Enzo life Inc. Blebbistatin (Cat #13013), Rho Kinase (ROCK) Inhibitor, Y-27632 (Cat #10005583) and Theaflavin 3,3’-digallate (TF-3) (Cat #25215) were purchased from Cayman Chemical Company.

5-Aza-2′-deoxycytidine (Aza) (Cat #A3565) was obtained from Sigma-Aldrich. Src inhibitor, Saracatinib (CAS #379231-04-6) was obtained from Santa Cruz Biotechnology. Anti-S1PR1 antibody (Cat #ASR-011) was acquired from Alomone Labs. Anti-Emerin (Cat #30853), anti-Src (Cat #2109), p-Src (Cat #12432), anti-Fak (Cat #3285T), anti-Lamin (Cat #86846S), anti- DNMT3a (Cat # 3598) antibodies from Cell Signaling Technology whereas, anti-actin (Cat #sc- 47778), anti-DNMT3b (Cat #sc-20704), anti-p-Tyr (PY99, Cat #sc-7020) and anti-p-Tyr (PY20, Cat # sc-508) were acquired from Santa Cruz Biotechnology. Anti-DNMT1 (Cat #39204) from Active Motif.

### Animals

Mice were approved by the Institutional Animal Care and Use Committee of the University of Illinois at Chicago. Four-five weeks old *EC-Fak^-/-^* and tdTomato-*EC-Fak^-/-^*mice received tamoxifen (80mg/kg, *i.p.*) for five consecutive days followed by a week of rest for drug wash out. We generated these mice by crossing *Fak^fl/fl^* with 5′Endo-SCL*-Fak-CreERT/*5′Endo- SCL*-CreER/Rosa-Tomato* lineage tracing mice line (Akhter et al., 2021; Balaji Ragunathrao et al., 2019; Liu et al., 2019). Tamoxifen (MilliporSigma #T5648) was prepared in corn oil (MilliporeSigma, #C8267) as described(Akhter et al., 2021). All experiments were performed on 6-8 weeks old mice of weight 20-25 gm. Sex-matched groups of male and female mice were used for these studies. No animals were excluded from analysis. We calculated sample size using software G Power based on pre-designed effect size between the groups based on Cohen’s principles. A power =0.80, significance level = 0.05 (Faul et al., 2007).

### Cell culture and transfection

Human pulmonary arterial endothelial cells (HPAE) obtained from Lonza Allendale, NJ, USA (Lonza #CC-2530) were used for all cell studies. Cells were cultured in a humidified atmosphere with 5% CO2 at 37°C in EGM-2 medium supplemented with growth factors (Lonza #CC-3162), 10% FBS and penicillin-streptomycin antibiotics. All studies were conducted on HPAE that were between passage 6-8. HPAE were transfected at passage 4-6 using Santa Cruz transfection reagent or Amaxa electroporation as described previously(Akhter et al., 2021). FAK was depleted using 5ʹ-AAACCAUCUUCAUCUUCCCUU -3ʹ (Dharmacon Inc)(Schmidt et al., 2013). MISSION^®^ shRNA (Sigma-Aldrich, Cat #SHCLND) was used to deplete KLF2. ON-TARGETplus Human SOX17 siRNA (Cat # J-013028-09-0002), ON-

TARGETplus Human FOXP1 siRNA (Cat # J-004256-17-0005) were acquired from Dharmacon Inc. Ambion™ Silencer™ Pre-Designed siRNA (Cat # 02665278) was used to deplete DNMT3a. In all experiments, control siRNA (siCtr) (ON-TARGETplus non-targeting pool (D-001810-10), Dharmacon Inc was used. Wild type and phosphodefective emerin mutant (Y74F/Y95F) cDNA were obtained from Guilluy Lab (Guilluy et al., 2014). HPAE were transfected with cDNA using FuGENE® HD Transfection Reagent (Promega #E2311) as described (Akhter et al., 2021).

### Atomic force microscopy

Young’s modulus (*E*) of cultured HPAEC was assessed by atomic force microscopy as described (Devine et al., 2020; Wong et al., 2020). Briefly, a silicon nitride cantilever with an 18° pyramid tip (MLCT, Bruker) was used for nanoindentation. The spring constant of the cantilever was determined from thermal fluctuations before each experiment. Indentation was performed under contact mode with force-distance 500 nm at 1 μm/s velocity, until trigger voltage (0.5 V) is reached, followed by retraction. Young’s modulus (*E*) was then calculated by fitting a force-indentation curve to the Hertzian model with Poisson’s ratio at 0.5.

### Confocal imaging of EC and lungs

HPAEC were rinsed three times with PBS and fixed with 2% paraformaldehyde. Cells were then incubated with 5% normal goat serum supplemented with 1% Bovine serum albumin in TBS (0.2M Tris base, 1.5M NaCl) for 2h at room temperature. After rinsing three times with PBS, cells were incubated with 1:50 anti-VE-Cadherin (Santa Cruz #sc- 515467) or anti-emerin (Cell Signaling #30853) antibody for 1h at room temperature following which cells were rinsed again and incubated with 1:250 dilution of Alexa-Fluor 594 secondary Donkey anti-rabbit antibody (ThermoFisher Scientific #A-11016) at room temperature for 1h. Isotype control primary antibodies (ThermoFisher Scientific, #31243 & 02-6102) were used as negative control to validate specificity of antibodies and to eliminate the background signal. DAPI (1:1000. ThermoFisher Scientific, #D-1306) was used for staining nuclei. The images were acquired with an inverted laser-scanning confocal microscope (LSM 880, Carl Zeiss Microscopy) using the Zeiss LSM software. Representative images shown in the figures were selected to most accurately match the quantitative analysis. Regions were selected randomly to avoid biasing.

Lungs were perfused with normal saline followed by administration of ice-cold 4% paraformaldehyde (PFA) solution for 2 h followed by equilibration with 30% sucrose solution overnight (Akhter et al., 2021). Lungs were then embedded in Optimal cutting temperature (OCT) compound and fast frozen at -20°C. Lungs were sectioned (8 to 10µm) and immunostained using anti-GFP antibody followed by vWF antibody. Antigen retrieval was performed in a citrate-based antigen unmasking solution (Vector Laboratories # H-3300) following manufacturer’s instructions with slide immersed in retrieval solution. Blocking was performed using 5% normal goat serum supplemented with 1% Bovine serum albumin in TBS (0.2M Tris base, 1.5M NaCl) for 2 h at room temperature.

### Stochastic optical reconstruction microscopy (STORM)

Control or FAK depleted EC were washed with PBS (X3), fixed with 4% paraformaldehyde for 10 min at room temperature followed by permeabilization in 0.1% Triton X-100 in PBS for 10 minutes at room temperature. After washing and blocking, cells were stained with anti-5mC antibody (1:1000 dilution) overnight followed by secondary antibody conjugated with fluorescent dyes Alexa Fluor 647 for 2 h at room temperature. STORM images were acquired using buffer A (0.5 mL 1M Tris (pH 8.0), 0.146 g NaCl, 50 ml water) and buffer B (2.5 mL 1M Tris (pH 8.0), 0.029 g NaCl, 5 g Glucose, 47.5 ml water). Imaging buffer (GLOX) was prepared as described(Xu et al., 2017). The images were acquired on GE OMX-SR Super-Resolution Microscope following manufacture’s protocol.

### Fluorescence activated cell sort

Lungs were minced and digested with 1mg/ml collagenase A (Roche #10103578001) for 30 min at 37°C after which digested tissue was passed through a 75μm cell strainer to obtain single-cell suspensions. Cells were stained with anti-CD31 and anti-CD45 antibodies and endothelial cell (CD31^+^CD45^-^) were sorted using Beckman Coulter cell sorter as described (Akhter et al., 2021).

### ATAC-seq

Endothelial cell (CD31^+^CD45^-^) sorted from control and tdTomato-*EC-Fak^-/-^* null mice were re-suspended in cold PBS after which chromatin was extracted and processed for Tn5- mediated tagmentation and adapter incorporation, according to the manufacturer’s protocol (Nextera DNA sample preparation kit, Illumina®) at 37 °C for 30 min. ATAC-seq was performed at Northwestern University Genomic Core, Chicago. Briefly, reduced-cycle amplification was carried out in the presence of compatible indexed sequencing adapters. The quality of the libraries was assessed by a DNA-based fluorometric assay (ThermoFisher Scientific^TM^) and automated capillary electrophoresis (Agilent Technologies, Inc.). Raw data have been deposited in gene expression omnibus archive (accession no. GSE207789).

### RNA-seq analysis

Endothelial cells (CD31^+^CD45^-^) were sorted from EC-FAK ^-/-^ null and control *FAK^fl/fl^* mice. RNA was isolated and quality control (QC) was performed using bioanalyzer. RNA-Seq analysis was performed in Genomic Core facility, Northwestern University, Chicago with 7-10 RIN (RNA Integrity Number). Bioinformatics analysis was performed as described (Akhter et al., 2021). Raw data have been deposited in gene expression omnibus archive (accession no. GSE212037).

### Quantitative real time PCR

Total RNA was isolated using Trizol reagent (ThermoFisher Scientific #15596026) and quantified using BioDrop DUO+ (Biochrom, UK). RNA (1µg) was reverse transcribed using High-Capacity RNA to cDNA Kit (Applied Biosystems #4368814) according to the manufacturer’s protocol. The cDNA products were assessed using quantitative Real Time PCR analysis and Fast SYBR™ Green Master Mix (Applied Biosystems # 4385612). Each measurement was carried out in duplicates using a CFX384 real Time on Applied Biosystems QuantStudio Real-Time PCR System. The PCR conditions were 95°C for 10 min followed by 40 cycles 95 °C for 15 sec and to 60 °C for 1 min. The quantitative real-time PCR data were analyzed by 2^-ΔΔCT^ method. The expression of each gene was normalized to GAPDH. The list of primers used has been provided in Table I as Online Supplementary data.

**Table 1.**
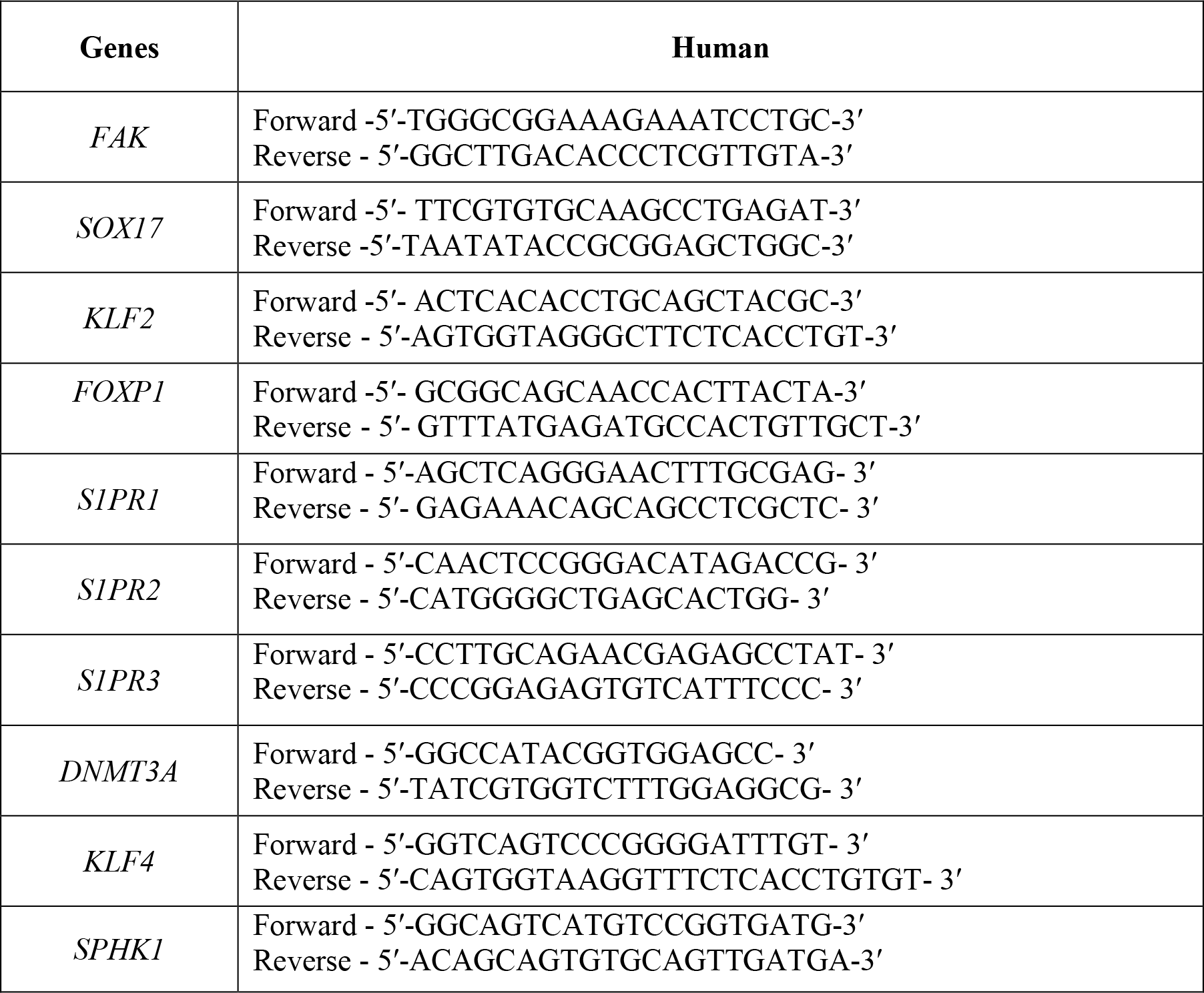
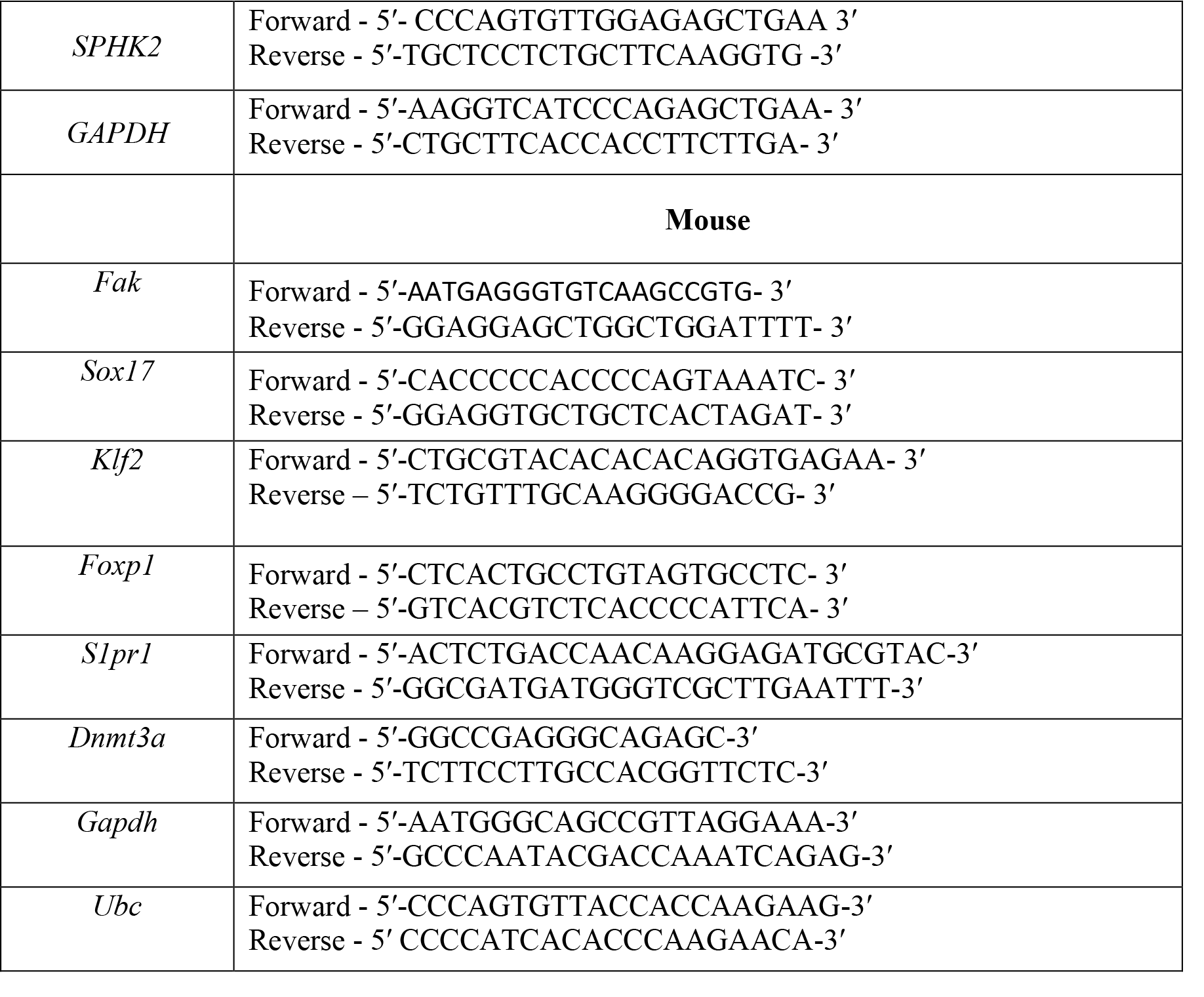
Primers list:

### Assessment of lung vascular injury

Lung wet-dry weight ratio were determined to quantify lung vascular injury as described previously (Akhter et al., 2021).

### Chromatin immunoprecipitation (*ChIP*) assay

Protein-DNA complex (100-120μg) was immunoprecipitated with the antibody against MEF2 as described previously (Akhter et al., 2021). Briefly, Protein-DNA complex (100–120μg) was immunoprecipitated with the anti-MEF2 antibody (Santa Cruz #sc-17785). DNA fragments were collected by phenol– chloroform–isoamyl alcohol extraction, followed by ethanol precipitation, and then resuspended in 14μl nuclease water for PCR. Promoter region of KLF2 targeted a 152bp fragment was quantified by Sybr green-based real time quantitative PCR (q-PCR) using ViiA7 (Applied Biosystem, Foster City, CA). Normal rabbit IgG was used as negative antibody control and DNA from the input (20–40 μg protein-DNA complexes) was used as an internal control. Primers used to amplify the MEF2 binding on KLF2 are forward 5ʹ-GCAGTCCGGGCTCCCGCAGTAG-3ʹ; reverse 5ʹ- CTTATAGGCGCGGCAGGCAC-3ʹ.

### Genomic DNA isolation and bisulfite sequencing

Genomic DNA was isolated from control or FAK depleted EC using the Qiagen Genomic DNA Kit (Qiagen # 69504). DNA (200 ng) was then processed for bisulfite conversion using the EpiGenTek Bisulfite Kit (#P-1054-050) according to the manufacturers’ instructions. The PCR sequence targeting *KLF2* promoter was performed under following conditions: denaturation at 98 °C for 60 s and 35 cycles each of 98 °C for 10 s, 55 °C for 30 s, and 72 °C for 45 s. The primer is listed in table 2. The PCR products were subcloned into the pGEM-T (pGEM-T Easy Vector System, Promega #A1360) using manufacture’s protocol, and individual clones were sequenced. The Sanger sequencing was performed at University of Illinois Genomic Core, UIC. The methylation status of the region was determined and analyzed with QUMA (quantification tool for methylation analysis) (http://quma.cdb.riken.jp/top/quma_main_j.html)(Kumaki et al., 2008).

**Table 2.**
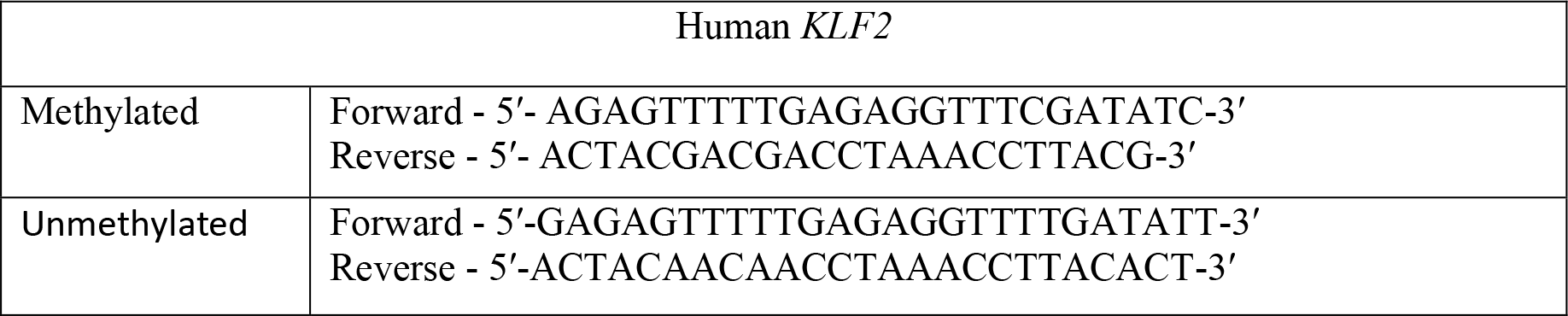
Methylation specific primers

### Methylation specific PCR

Since methylation at CpG sites on the promoter of a gene down regulates its transcription, we quantified DNA methylation at the promoter region of KLF2 by bisulfite conversion of the DNA. Bisulfite reaction was performed using EpiTect Plus Lyse All Bisulfite Kit (Qiagen #59124), and after sodium bisulfite conversion, the DNA was applied to an EpiTect spin column and washed to remove traces of sodium bisulfite. The converted DNA was eluted with nuclease-free water. PCR reaction was performed for methylated (M) and un- methylated (U) regions using 2 uL of 5X PCR reaction buffer, 0.2 mM of dNTPs (Promega, Madison, WI), 0.2 uM of each forward and reverse primers, GoTaq DNA polymerase, and 1 uL converted DNA. The primers for methylated and unmethylated sequences listed in Table 2, were designed using online MethPrimer(Li and Dahiya, 2002). Thermal cycling conditions included 94°C for 5 minutes, 35 cycles of 94°C for 1 minute, 55°C for 30 seconds, 72°C for 1 minute, and final extension at 72°C for 10 minutes. The PCR products were analyzed on 2% agarose gel and the intensity of methylated to unmethylated bands was quantified.

### DNA methyltransferase activity measurement

Nuclear fraction was isolated from culture HPAE using NE-PER™ Nuclear and Cytoplasmic Extraction Reagents (Thermofisher Scientific #78833) as per the manufacture’s protocol. The nuclear lysates were used to assess the DNMT activity following manufacture’s method of EpiQuik DNA Methyltransferase Activity Assay Kit (EpiGenTek # P-3001-1).

### Western blotting

HPAE were lysed using radioimmunoprecipitation assay buffer (RIPA buffer containing 10 mM Tris-HCl pH 8.0, 1 mM EDTA, 0.5 mM EGTA, 1% Triton X-100, 0.1% sodium deoxycholate, 0.1% SDS, 140 mM NaCl and 1 mM phenylmethylsulfonyl fluoride (PMSF). Western blot experiments were performed using indicated antibodies as described previously (Akhter et al., 2021).

### Immunoprecipitation

For immunoprecipitation analysis, cells were washed with ice cold PBS and immediately processed for sub-cellular fractionation as described (Rajput et al., 2013). Sub cellular nuclear lysates were incubated with anti-Phospho tyrosine (P99 and P20, 1:250) antibodies overnight followed by addition of agarose beads to pull down the immune complexes (Rajput et al., 2013). Proteins were separated by SDS-PAGE and immunoblotted using indicated antibodies. Dilution for each primary antibody used in the study were as follows Emerin (1:1000), Lamin A (1:1000), Phospho tyrosine (P99 and P20, 1:250). Membranes were then incubated with respective secondary antibodies anti-mouse or anti rabbit (1:10000) for 2 h following which the bands were visualized using imager or autoradiographic films and chemiluminescent western blotting detection substrate.

### Trans-endothelial electrical resistance (TEER)

HPAE were seeded on gelatin coated gold-plated eight-well electrodes (8W standard Array; 8W10E+) (Applied Biosciences, Carlsbad, CA). Cells were transfected with indicated siRNAs for 48 h. The smaller electrode and larger counter electrode were then connected to a phase-sensitive lock-in amplifier to monitor the voltage. A constant current of 1 μA was supplied by a 1 V, 4000 Hz AC signal connected serially to a 1 MΩ resistor between the smaller electrode and the larger counter electrode. EC monolayers were incubated in serum-free medium for 2 h following which S1P was added to assess dynamic change in TEER (Akhter et al., 2021).

### Liposome-mediated *in-vivo* gene delivery

Liposomes were prepared using a mixture of chloroform, Dimethyl Dioctadecyl Ammonium Bromide and cholesterol as described previously (Tauseef et al., 2012). Briefly, chloroform was evaporated from the mixture using a rotavapor system at a speed of 105 rpm for 15-20 min at 37°C to make a lipid layer. Lipid layer was extracted by sonicating the solution for 1 h at 42°C in presence of 5% glucose. The liposomes were filtered through a 0.45-micron filter and subsequently cDNA was added. The cDNA loaded liposomes were administered i.v into the mouse. The lungs were harvested for edema measurement after 24 h.

### Polyethylene glycol hydrogel Preparation

The soft and stiff hydrogels were formulated as described (Lenzini et al., 2021). Briefly, these hydrogels were prepared under the exposure of ultraviolet light (365 nm) by adding varying concentrations of lithium phenyl(2,4,6- trimethylbenzoyl) phosphinate (TCI Chemicals) to the solution of PEG-DA Mn 700, 10 mM sodium L-ascorbate, 4 mM tris(2-carboxyethyl) phosphine, 1× phosphate-buffered saline. Cys- RGD peptide (sequence CGGGGRGDSP) at a concentration of 0.16 mM was added to the pre-hydrogel solution for the attachment of the endothelial cells. Increased amount of LAP promotes to achieve improved mechanical strength in terms of Young’s modulus.

### Statistical analysis

Data are presented as the mean ± SEM. Wherever only two groups are compared statistical differences were assessed using unpaired two-tailed Student’s t test. Statistical significance among three groups or over was determined using one-way analysis of variance (ANOVA) followed by post hoc Tukey’s multiple comparisons test. The data were analyzed using GraphPad Prism version 8.0, and P < 0.05 was considered to indicate a statistically significant difference.

## Acknowledgements

This work was supported by the National Institutes of Health, USA (grants HL060678, HL137169 and HL084153). MZA was supported by American Heart Association Postdoctoral Fellowship (AHA Award-19POST34450241). We thank Dr. Christophe Guilluy, Institute for Advanced Biosciences, Centre de recherche UGA–INSERM, France for generously providing us the WT emerin and phosphodefective mutants cDNAs. We would like to acknowledge Lakshmi Yalagala for skillful performance of the mouse genotyping.

## Author contributions

M.Z.A. and D.M. designed the experiments. M.Z.A., P.Y., M.T., M.A., F.H., S.D., V.V., J.C.J., N.S., S.L., G.Z., and J.W.S. performed the experiments and analyzed data.

J.L and M.K.J provided reagents. M.Z.A. and D.M. wrote the manuscript. M.A., S.D., V.V., J.C.J., N.A., G.Z., J.L, M.K.J and J.W.S commented on the manuscript.

## Supplementary Figures

Figure 1: **Role of FAK in altering mechanotransduction and chromatin accessibility. A-B**, Immunoblot **(A)** shows FAK depletion in EC while **B** shows densitometric analysis of FAK expression. Actin was used as loading control. Experiments were performed three times independently. **C**, EC were transfected with vector, WT-RhoA, dominant-negative (T19N), or constitutively active RhoA (Q63L-RhoA) cDNA constructs for 24h, and intracellular tension was measured by AFM in a single GFP-expressing EC. N= 40 cells per group pooled from three independent experiments. **D**, ATAC-seq heatmap showing genome-wide differential open chromatin peaks in FAK^+^ EC versus FAK^-^ EC. Total number of peaks 52,500. **E**, Read-count frequency of genomic regions from EC sorted from control or FAK-null lungs. **F**, mRNA expression of *KLF2*, *FOXP1*, *SOX17* and *FAK* in control and FAK-depleted human EC using qPCR. GAPDH was used as the house keeping gene (n=3). Data plotted in **B, C** and **F** are given as mean ± SEM. Statistical significance was assessed by one-way ANOVA followed by Post hoc Tukey’s test in C, and unpaired t-test was used for B and F. \*\*\**P*< 0.001, \*\**P*< 0.01, relative to GFP and RhoA transduced cells (C) and siCtr (B, F).

Figure 2: **KLF2 regulates endothelial barrier function downstream of FAK. A-C,** mRNA expression of *KLF2*, *FOXP1* and *SOX17* measured by qPCR using GAPDH as internal control (n=3). **D**, mRNA expression of *KLF2* in FAK depleted EC transducing KLF2 cDNA. GAPDH was used as an internal control (n=3). **E,** TEER was measured as described in Fig. 1G in FAK- depleted EC after transducing vector, KLF2, or SOX17 cDNA post 48 h of FAK depletion (*n* = 8/group). Basal barrier function was assessed 72 h post-transfection. **F,** mRNA expression of *SOX17* in FAK depleted EC transducing SOX17 cDNA. GAPDH was used as an internal control (n=3). **G,** Micrograph of a lung section receiving GFP tagged KLF2 cDNA. Anti-vWF antibody was used to mark the endothelium. Scale bar, 20 µm. Experiment was repeated three times independently. Data plotted in **A, B, C, D, E** and **F** are given as mean ± SEM. Statistical significance was assessed by one-way ANOVA followed by Post hoc Tukey’s test in **D, E** and **F** and unpaired t-test in **A, B** and **C.** \*\*\*\**P*< 0.0001, \*\*\**P*< 0.001, \*\**P*< 0.01 relative to shCtr (A), siCtr (B, C), siCtr+vector (D, E, F).

Figure 3: **DNMT3a regulates *KLF2* expression. A,** DNA-PAGE of ChIP qPCR amplified products showing MEF2 binding to *KLF2* in control but not in FAK-depleted EC. IgG was used as control. **B,** Representative immunoblots of indicated proteins in three independent experiments. Actin was used as loading control. Lysates were prepared from lungs of *Fak^fl/fl^* and *EC-Fak^-/-^* mice (n=3). **C,** Corresponding densitometric analysis of above blots. **D,** mRNA expression of indicated genes in control, FAK depleted, and FAK + DNMT3a-depleted EC. GAPDH was used an internal control (n=3). **E-F,** WT mice were challenged with LPS (10 mg/kg, *i.p*) and after 8 h lungs were harvested. Lung sections were immunostained with anti-FAK antibody and anti-vWF antibody after which images were acquired using confocal imaging. **E** shows a representative micrograph and **F**, quantification of FAK expression in vessels (n=3). Data plotted in **C, D** and **F** are given as mean ± SEM. Statistical significance was assessed by one-way ANOVA followed by Post hoc Tukey’s test in **D** and unpaired t-test was used in **C and F**. \*\*\**P*< 0.001, \*\**P*< 0.01 relative to *Fak^fl/fl^* (C) siCtr (D), No LPS (F).

Figure 4: **Inhibition of DNMT3a activity restores KLF2 and edema. A-B,** mRNA expression of *KLF2* in FAK-depleted EC treated for 4 h with the AZA (500 nM, A) or TF-3 (1 µM, B). GAPDH was used an internal control (n=3). **C,** *Fak^fl/fl^* and *EC-Fak^-/-^* mice were injected with AZA (0.5 mg/kg, *i.v.*) or TF-3 (1mg/kg, *i.v*). Lungs were harvested after 24 h and mRNA expression of *KLF2* was assessed by qPCR. GAPDH was used as internal control (n=3). **D,** A representative immunoblot show DNMT3a and FAK deletion in lung lysates prepared from double knockout and control mice. Experiments were repeated two times independently. **E,** Lung edema was assessed by measuring lung weight-dry weight ratio in *Fak^fl/fl^* and *EC-Fak^-/-^* after without or with AZA (0.5mg/kg, i.v) (n=6 mice/gp). **F,** DNMT activity was assessed in EC cultured on soft and stiff PEG gels for 72 h as described in 2G. Experiments were repeated two times independently. Data plotted in **A, B, C, E** and **F** are given as mean ± SEM. Statistical significance was assessed by one- way ANOVA followed by Post hoc Tukey’s test in A, B, C and E. \*\*\**P*< 0.001, \*\**P*< 0.01, \**P*< 0.05 relative to *Fak^fl/fl^* (C, E), siCtr+AZA (A), siCtr+TF3 (B), ###*P*< 0.001 relative to *EC- Fak^-/-^* (E).

Figure 5: **Effect of EC-FAK deletion on EC transcriptome**. **A-B,** Representative blot **(A)** and densitometry **(B)** of S1PR1 expression as fold change over actin (n=3). **C,** Expression of indicated genes in control versus FAK-depleted EC. GAPDH was used as an internal control (n=3). **D,** Quantification of TEER in FAK-depleted EC before and after stimulation with 1 µM S1P (n=3 wells/group). **E-F,** Immunoblot of S1PR1 expression in lung lysates from mice receiving S1PR1 cDNA **(E)**. Actin was used as loading control. **F,** densitometric analysis of the above blots. (n=3). **G,** Quantification of TEER. EC plated on TEER electrodes were transfected with control or KLF2 shRNA. After 48 h, cells were stimulated with S1P (1 µM) and TEER was measured as in **2E** (*n* = 8wells/group). Data plotted in **B, C, D, F** and **G** are given as mean ± SEM. Statistical significance was assessed by one-way ANOVA followed by Post hoc Tukey’s test in D and F and unpaired t- test in B, C and G. \*\*\*\**P*< 0.0001, \*\*\**P*< 0.001, \*\**P*< 0.01 relative to siCtr (B, C, D), *EC-Fak^-/-^*receiving vector (F), shCtr (G).

Figure 6: **Inhibition of emerin activity restores S1PR1 expression. A-B,** Representative micrograph shows emerin localization in nuclei in EC cultured on soft versus stiff matrix. The EC were stained with anti-emerin antibody and an appropriate fluorescent secondary antibody. Nuclei were labelled using DAPI to assess emerin localization. Scale bar: 20 µm. Data were acquired from two independent experiments. **B,** Quantitation of emerin peripheral reorganization as described in Fig. 6A. **C,** Plot shows changes in cSrc phosphorylation in FAK-depleted EC after without or with treatment with the indicated inhibitors. cSrc was used as a loading control (n=3). **D,** Representative immunoblot of FAK and emerin in FAK-depleted EC transduced with WT- or a phosphodefective emerin mutant (Y74-Y95FF) from two independent experiments. Actin was used as a loading control. **E-F,** mRNA expression of *S1PR1* in FAK-depleted EC treated with the AZA (500 nM) for 4h **(E)**. **F,** in lung of WT mice injected with of TF-3 administration (1 mg/kg, i.v) the mice after 1 h, received LPS (10 mg/kg, *i.p*) as described in Fig. 3A. Data plotted in **B, C, E** and **F** are given as mean ± SEM. Statistical significance was assessed by one-way ANOVA followed by Post hoc Tukey’s test in **C, E and F** and unpaired t-test was used in **B**. \*\**P*< 0.01, \**P*< 0.05 relative to soft (B), siCtr (C), siCtr +AZA (E), No LPS (F) and # *P*< 0.05 LPS (F).

## References

1. Akhter, M.Z., Chandra Joshi, J., Balaji Ragunathrao, V.A., Maienschein-Cline, M., Proia, R.L., Malik, A.B., and Mehta, D. (2021). Programming to S1PR1(+) Endothelial Cells Promotes Restoration of Vascular Integrity. Circ Res 129, 221–236.

2. Atkins, G.B., and Jain, M.K. (2007). Role of Kruppel-like transcription factors in endothelial biology. Circ Res 100, 1686–1695.

3. Balaji Ragunathrao, V.A., Anwar, M., Akhter, M.Z., Chavez, A., Mao, Y., Natarajan, V., Lakshmikanthan, S., Chrzanowska-Wodnicka, M., Dudek, A.Z., Claesson-Welsh, L., et al. (2019). Sphingosine-1-Phosphate Receptor 1 Activity Promotes Tumor Growth by Amplifying VEGF- VEGFR2 Angiogenic Signaling. Cell Rep 29, 3472–3487 e3474.

4. Bastounis, E.E., Yeh, Y.T., and Theriot, J.A. (2019). Subendothelial stiffness alters endothelial cell traction force generation while exerting a minimal effect on the transcriptome. Sci Rep 9, 18209.

5. Bhattacharya, J., and Matthay, M.A. (2013). Regulation and repair of the alveolar-capillary barrier in acute lung injury. Annu Rev Physiol 75, 593–615.

6. Birukova, A.A., Arce, F.T., Moldobaeva, N., Dudek, S.M., Garcia, J.G., Lal, R., and Birukov, K.G. (2009). Endothelial permeability is controlled by spatially defined cytoskeletal mechanics: atomic force microscopy force mapping of pulmonary endothelial monolayer. Nanomedicine 5, 30–41.

7. Birukova, A.A., Tian, X., Cokic, I., Beckham, Y., Gardel, M.L., and Birukov, K.G. (2013). Endothelial barrier disruption and recovery is controlled by substrate stiffness. Microvasc Res 87, 50–57.

8. Braren, R., Hu, H., Kim, Y.H., Beggs, H.E., Reichardt, L.F., and Wang, R. (2006). Endothelial FAK is essential for vascular network stability, cell survival, and lamellipodial formation. J Cell Biol 172, 151–162.

9. Christman, J.K. (2002). 5-Azacytidine and 5-aza-2’-deoxycytidine as inhibitors of DNA methylation: mechanistic studies and their implications for cancer therapy. Oncogene 21, 5483–5495.

10. Daneshjou, N., Sieracki, N., van Nieuw Amerongen, G.P., Conway, D.E., Schwartz, M.A., Komarova, Y.A., and Malik, A.B. (2015). Rac1 functions as a reversible tension modulator to stabilize VE-cadherin trans-interaction. J Cell Biol 208, 23–32.

11. Delgadillo, L.F., Marsh, G.A., and Waugh, R.E. (2020). Endothelial Glycocalyx Layer Properties and Its Ability to Limit Leukocyte Adhesion. Biophys J 118, 1564–1575.

12. Denis, H., Ndlovu, M.N., and Fuks, F. (2011). Regulation of mammalian DNA methyltransferases: a route to new mechanisms. EMBO Rep 12, 647–656.

13. Devine, D., Vijayakumar, V., Wong, S.W., Lenzini, S., Newman, P., and Shin, J.W. (2020). Hydrogel Micropost Arrays with Single Post Tunability to Study Cell Volume and Mechanotransduction. Adv Biosyst 4, e2000012.

14. Discher, D.E., Janmey, P., and Wang, Y.L. (2005). Tissue cells feel and respond to the stiffness of their substrate. Science 310, 1139–1143.

15. Even-Ram, S., Doyle, A.D., Conti, M.A., Matsumoto, K., Adelstein, R.S., and Yamada, K.M. (2007). Myosin IIA regulates cell motility and actomyosin-microtubule crosstalk. Nat Cell Biol 9, 299–309.

16. Faul, F., Erdfelder, E., Lang, A.G., and Buchner, A. (2007). G*Power 3: a flexible statistical power analysis program for the social, behavioral, and biomedical sciences. Behav Res Methods 39, 175–191.

17. Garcia, J.G., Liu, F., Verin, A.D., Birukova, A., Dechert, M.A., Gerthoffer, W.T., Bamberg, J.R., and English, D. (2001). Sphingosine 1-phosphate promotes endothelial cell barrier integrity by Edg-dependent cytoskeletal rearrangement. J Clin Invest 108, 689–701.

18. Guilluy, C., and Burridge, K. (2015). Nuclear mechanotransduction: forcing the nucleus to respond. Nucleus 6, 19–22.

19. Guilluy, C., Osborne, L.D., Van Landeghem, L., Sharek, L., Superfine, R., Garcia-Mata, R., and Burridge, K. (2014). Isolated nuclei adapt to force and reveal a mechanotransduction pathway in the nucleus. Nat Cell Biol 16, 376–381.

20. Guo, X., Wang, L., Li, J., Ding, Z., Xiao, J., Yin, X., He, S., Shi, P., Dong, L., Li, G., et al. (2015). Structural insight into autoinhibition and histone H3-induced activation of DNMT3A. Nature 517, 640–644.

21. Hariri, L., and Hardin, C.C. (2020). Covid-19, Angiogenesis, and ARDS Endotypes. N Engl J Med 383, 182–183.

22. Higuchi, M., Ishiyama, K., Maruoka, M., Kanamori, R., Takaori-Kondo, A., and Watanabe, N. (2021). Paradoxical activation of c-Src as a drug-resistant mechanism. Cell Rep 34, 108876.

23. Hoffman, B.D., Grashoff, C., and Schwartz, M.A. (2011). Dynamic molecular processes mediate cellular mechanotransduction. Nature 475, 316–323.

24. Holinstat, M., Knezevic, N., Broman, M., Samarel, A.M., Malik, A.B., and Mehta, D. (2006). Suppression of RhoA activity by focal adhesion kinase-induced activation of p190RhoGAP: role in regulation of endothelial permeability. J Biol Chem 281, 2296–2305.

25. Ingber, D.E. (2002). Mechanical signaling and the cellular response to extracellular matrix in angiogenesis and cardiovascular physiology. Circ Res 91, 877–887.

26. Knezevic, N., Tauseef, M., Thennes, T., and Mehta, D. (2009). The G protein betagamma subunit mediates reannealing of adherens junctions to reverse endothelial permeability increase by thrombin. J Exp Med 206, 2761–2777.

27. Kumaki, Y., Oda, M., and Okano, M. (2008). QUMA: quantification tool for methylation analysis. Nucleic Acids Res 36, W170–175.

28. Kumar, A., Kumar, S., Vikram, A., Hoffman, T.A., Naqvi, A., Lewarchik, C.M., Kim, Y.R., and Irani, K. (2013). Histone and DNA methylation-mediated epigenetic downregulation of endothelial Kruppel-like factor 2 by low-density lipoprotein cholesterol. Arterioscler Thromb Vasc Biol 33, 1936–1942.

29. Lathen, C., Zhang, Y., Chow, J., Singh, M., Lin, G., Nigam, V., Ashraf, Y.A., Yuan, J.X., Robbins, I.M., and Thistlethwaite, P.A. (2014). ERG-APLNR axis controls pulmonary venule endothelial proliferation in pulmonary veno-occlusive disease. Circulation 130, 1179–1191.

30. Lenzini, S., Debnath, K., Joshi, J.C., Wong, S.W., Srivastava, K., Geng, X., Cho, I.S., Song, A., Bargi, R., Lee, J.C., et al. (2021). Cell-Matrix Interactions Regulate Functional Extracellular Vesicle Secretion from Mesenchymal Stromal Cells. ACS Nano.

31. Li, L.C., and Dahiya, R. (2002). MethPrimer: designing primers for methylation PCRs. Bioinformatics 18, 1427–1431.

32. Li, S., Butler, P., Wang, Y., Hu, Y., Han, D.C., Usami, S., Guan, J.L., and Chien, S. (2002). The role of the dynamics of focal adhesion kinase in the mechanotaxis of endothelial cells. Proc Natl Acad Sci U S A 99, 3546–3551.

33. Li, Y., and Tollefsbol, T.O. (2011). DNA methylation detection: bisulfite genomic sequencing analysis. Methods Mol Biol 791, 11–21.

34. Liu, M., Zhang, L., Marsboom, G., Jambusaria, A., Xiong, S., Toth, P.T., Benevolenskaya, E.V., Rehman, J., and Malik, A.B. (2019). Sox17 is required for endothelial regeneration following inflammation-induced vascular injury. Nat Commun 10, 2126.

35. Matthay, M.A., and Zemans, R.L. (2011). The acute respiratory distress syndrome: pathogenesis and treatment. Annu Rev Pathol 6, 147–163.

36. McVerry, B.J., Peng, X., Hassoun, P.M., Sammani, S., Simon, B.A., and Garcia, J.G. (2004). Sphingosine 1-phosphate reduces vascular leak in murine and canine models of acute lung injury. Am J Respir Crit Care Med 170, 987–993.

37. Mehta, D., and Malik, A.B. (2006). Signaling mechanisms regulating endothelial permeability. Physiol Rev 86, 279–367.

38. Mehta, D., Tiruppathi, C., Sandoval, R., Minshall, R.D., Holinstat, M., and Malik, A.B. (2002). Modulatory role of focal adhesion kinase in regulating human pulmonary arterial endothelial barrier function. J Physiol 539, 779–789.

39. Parmar, K.M., Larman, H.B., Dai, G., Zhang, Y., Wang, E.T., Moorthy, S.N., Kratz, J.R., Lin, Z., Jain, M.K., Gimbrone, M.A., Jr., et al. (2006). Integration of flow-dependent endothelial phenotypes by Kruppel-like factor 2. J Clin Invest 116, 49–58.

40. Parsons, J.T. (2003). Focal adhesion kinase: the first ten years. J Cell Sci 116, 1409–1416.

41. Quadri, S.K. (2012). Cross talk between focal adhesion kinase and cadherins: role in regulating endothelial barrier function. Microvasc Res 83, 3–11.

42. Rajput, C., Kini, V., Smith, M., Yazbeck, P., Chavez, A., Schmidt, T., Zhang, W., Knezevic, N., Komarova, Y., and Mehta, D. (2013). Neural Wiskott-Aldrich syndrome protein (N-WASP)-mediated p120-catenin interaction with Arp2-Actin complex stabilizes endothelial adherens junctions. J Biol Chem 288, 4241–4250.

43. Russell-Hallinan, A., Watson, C.J., O’Dwyer, D., Grieve, D.J., and O’Neill, K.M. (2021). Epigenetic Regulation of Endothelial Cell Function by Nucleic Acid Methylation in Cardiac Homeostasis and Disease. Cardiovasc Drugs Ther 35, 1025–1044.

44. Schmidt, T.T., Tauseef, M., Yue, L., Bonini, M.G., Gothert, J., Shen, T.L., Guan, J.L., Predescu, S., Sadikot, R., and Mehta, D. (2013). Conditional deletion of FAK in mice endothelium disrupts lung vascular barrier function due to destabilization of RhoA and Rac1 activities. Am J Physiol Lung Cell Mol Physiol 305, L291–300.

45. Sei, Y.J., Ahn, S.I., Virtue, T., Kim, T., and Kim, Y. (2017). Detection of frequency-dependent endothelial response to oscillatory shear stress using a microfluidic transcellular monitor. Sci Rep 7, 10019.

46. SenBanerjee, S., Lin, Z., Atkins, G.B., Greif, D.M., Rao, R.M., Kumar, A., Feinberg, M.W., Chen, Z., Simon, D.I., Luscinskas, F.W., et al. (2004). KLF2 Is a novel transcriptional regulator of endothelial proinflammatory activation. J Exp Med 199, 1305–1315.

47. Seong, J., Tajik, A., Sun, J., Guan, J.L., Humphries, M.J., Craig, S.E., Shekaran, A., Garcia, A.J., Lu, S., Lin, M.Z., et al. (2013). Distinct biophysical mechanisms of focal adhesion kinase mechanoactivation by different extracellular matrix proteins. Proc Natl Acad Sci U S A 110, 19372–19377.

48. Sieg, D.J., Ilic, D., Jones, K.C., Damsky, C.H., Hunter, T., and Schlaepfer, D.D. (1998). Pyk2 and Src-family protein-tyrosine kinases compensate for the loss of FAK in fibronectin-stimulated signaling events but Pyk2 does not fully function to enhance FAK- cell migration. EMBO J 17, 5933–5947.

49. Skon, C.N., Lee, J.Y., Anderson, K.G., Masopust, D., Hogquist, K.A., and Jameson, S.C. (2013). Transcriptional downregulation of S1pr1 is required for the establishment of resident memory CD8+ T cells. Nat Immunol 14, 1285–1293.

50. Tauseef, M., Kini, V., Knezevic, N., Brannan, M., Ramchandaran, R., Fyrst, H., Saba, J., Vogel, S.M., Malik, A.B., and Mehta, D. (2008). Activation of sphingosine kinase-1 reverses the increase in lung vascular permeability through sphingosine-1-phosphate receptor signaling in endothelial cells. Circ Res 103, 1164–1172.

51. Tauseef, M., Knezevic, N., Chava, K.R., Smith, M., Sukriti, S., Gianaris, N., Obukhov, A.G., Vogel, S.M., Schraufnagel, D.E., Dietrich, A., et al. (2012). TLR4 activation of TRPC6-dependent calcium signaling mediates endotoxin-induced lung vascular permeability and inflammation. J Exp Med 209, 1953–1968.

52. Tifft, K.E., Bradbury, K.A., and Wilson, K.L. (2009). Tyrosine phosphorylation of nuclear-membrane protein emerin by Src, Abl and other kinases. J Cell Sci 122, 3780–3790.

53. Tomakidi, P., Schulz, S., Proksch, S., Weber, W., and Steinberg, T. (2014). Focal adhesion kinase (FAK) perspectives in mechanobiology: implications for cell behaviour. Cell Tissue Res 357, 515–526.

54. Tran, K.A., Zhang, X., Predescu, D., Huang, X., Machado, R.F., Gothert, J.R., Malik, A.B., Valyi-Nagy, T., and Zhao, Y.Y. (2016). Endothelial beta-Catenin Signaling Is Required for Maintaining Adult Blood-Brain Barrier Integrity and Central Nervous System Homeostasis. Circulation 133, 177–186.

55. Uhler, C., and Shivashankar, G.V. (2017). Regulation of genome organization and gene expression by nuclear mechanotransduction. Nat Rev Mol Cell Biol 18, 717–727.

56. Wang, N., Tytell, J.D., and Ingber, D.E. (2009). Mechanotransduction at a distance: mechanically coupling the extracellular matrix with the nucleus. Nat Rev Mol Cell Biol 10, 75–82.

57. Watanabe, K., Ueno, M., Kamiya, D., Nishiyama, A., Matsumura, M., Wataya, T., Takahashi, J.B., Nishikawa, S., Nishikawa, S., Muguruma, K., et al. (2007). A ROCK inhibitor permits survival of dissociated human embryonic stem cells. Nat Biotechnol 25, 681–686.

58. Wong, S.W., Lenzini, S., Bargi, R., Feng, Z., Macaraniag, C., Lee, J.C., Peng, Z., and Shin, J.W. (2020). Controlled Deposition of 3D Matrices to Direct Single Cell Functions. Adv Sci (Weinh) 7, 2001066.

59. Xia, S., Yu, W., Menden, H., Younger, S.T., and Sampath, V. (2021). FOXC2 Autoregulates Its Expression in the Pulmonary Endothelium After Endotoxin Stimulation in a Histone Acetylation-Dependent Manner. Front Cell Dev Biol 9, 657662.

60. Xu, J., Ma, H., and Liu, Y. (2017). Stochastic Optical Reconstruction Microscopy (STORM). Curr Protoc Cytom 81, 12 46 11-12 46 27.

61. Ying, L., Alvira, C.M., and Cornfield, D.N. (2018). Developmental differences in focal adhesion kinase expression modulate pulmonary endothelial barrier function in response to inflammation. Am J Physiol Lung Cell Mol Physiol 315, L66–L77.

62. You, D., Nilsson, E., Tenen, D.E., Lyubetskaya, A., Lo, J.C., Jiang, R., Deng, J., Dawes, B.A., Vaag, A., Ling, C., et al. (2017). Dnmt3a is an epigenetic mediator of adipose insulin resistance. Elife 6.

63. Zhou, J., Aponte-Santamaria, C., Sturm, S., Bullerjahn, J.T., Bronowska, A., and Grater, F. (2015). Mechanism of Focal Adhesion Kinase Mechanosensing. PLoS Comput Biol 11, e1004593.

