## Supplementary figures and images for "Tension sensing by FAK governs nuclear mechanotransduction, endothelial transcriptome and fate"

### Supplementary Figures

**Supplementary 1**

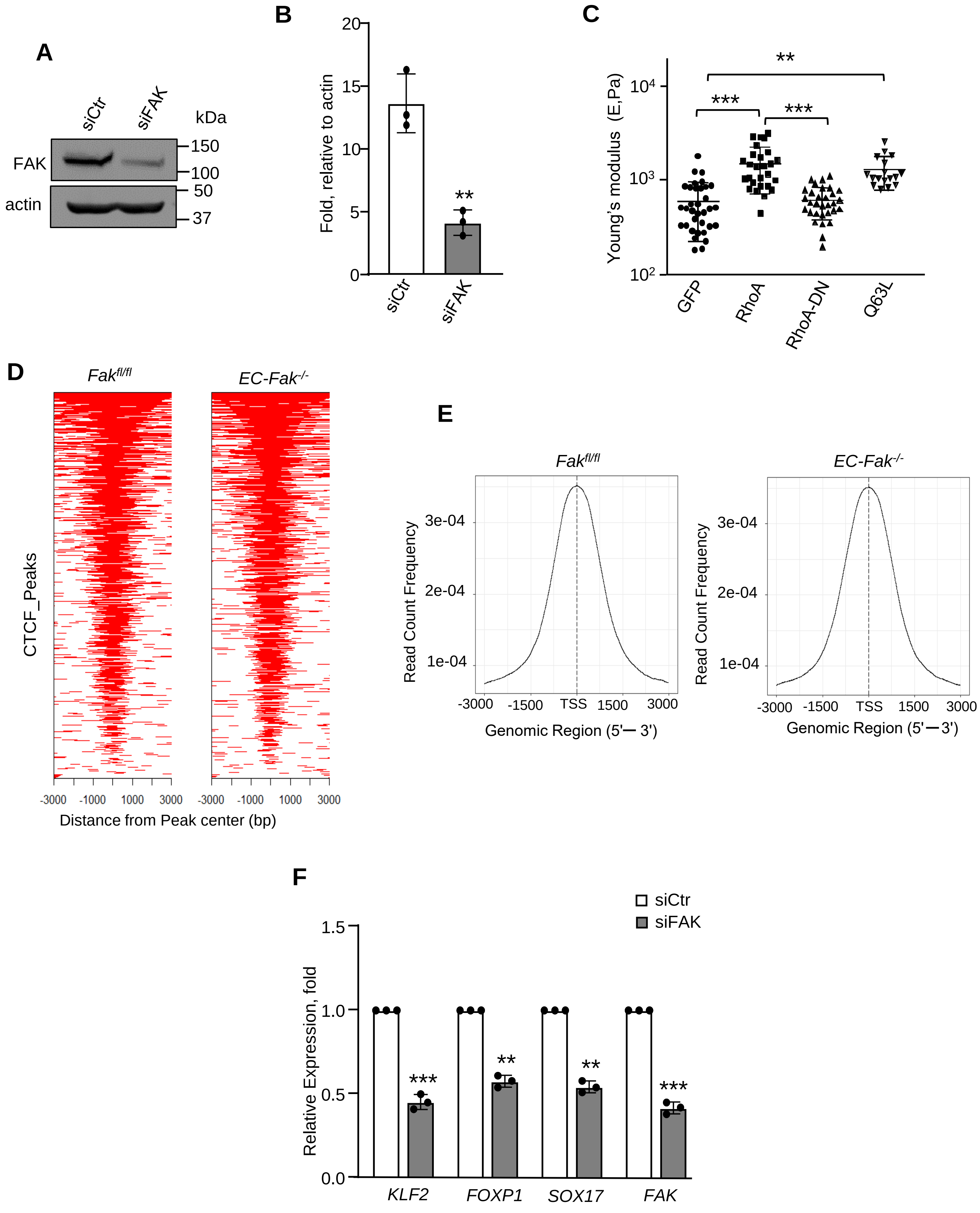

##### Supplementary 2

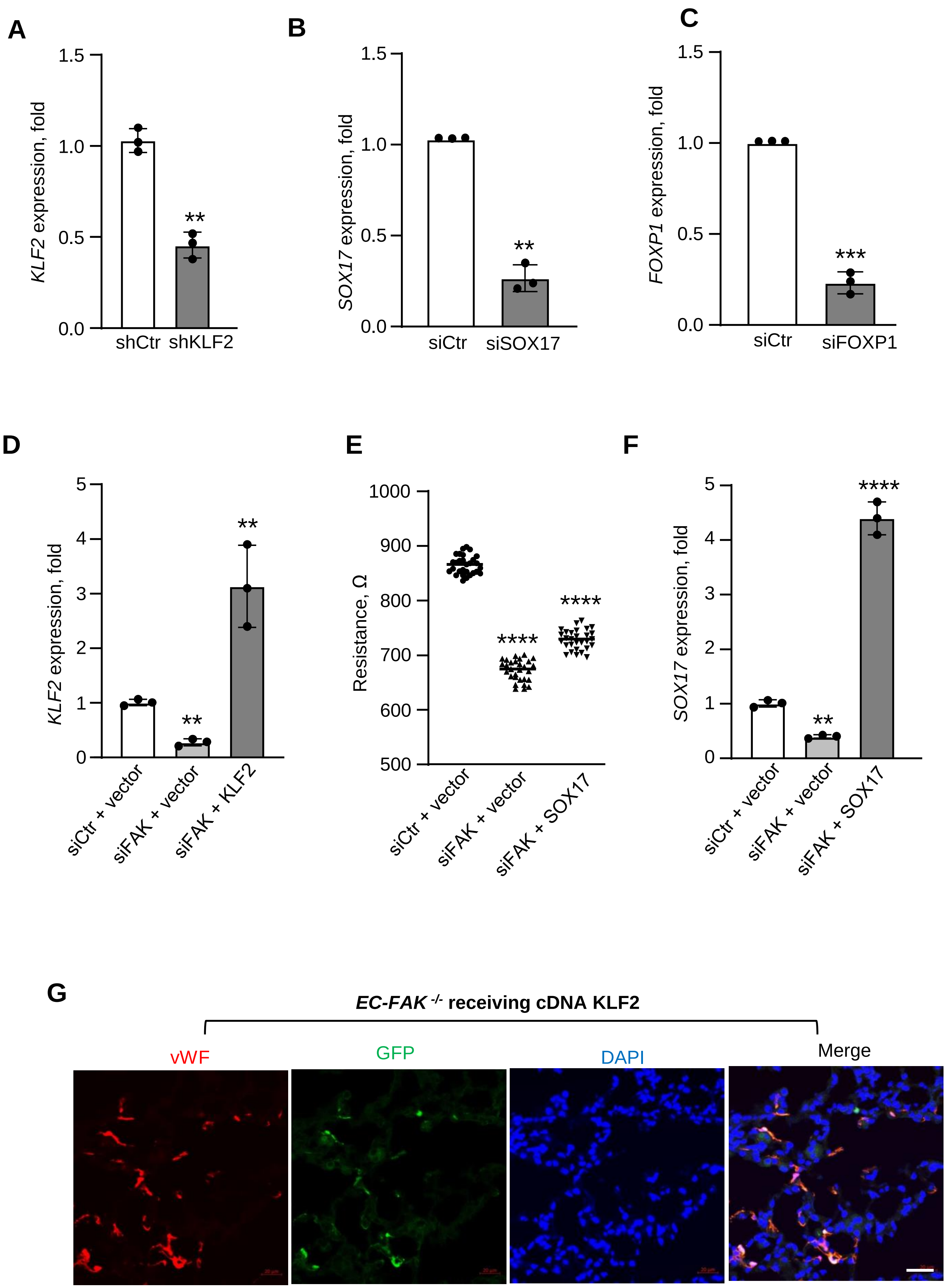

#### Supplementary 3

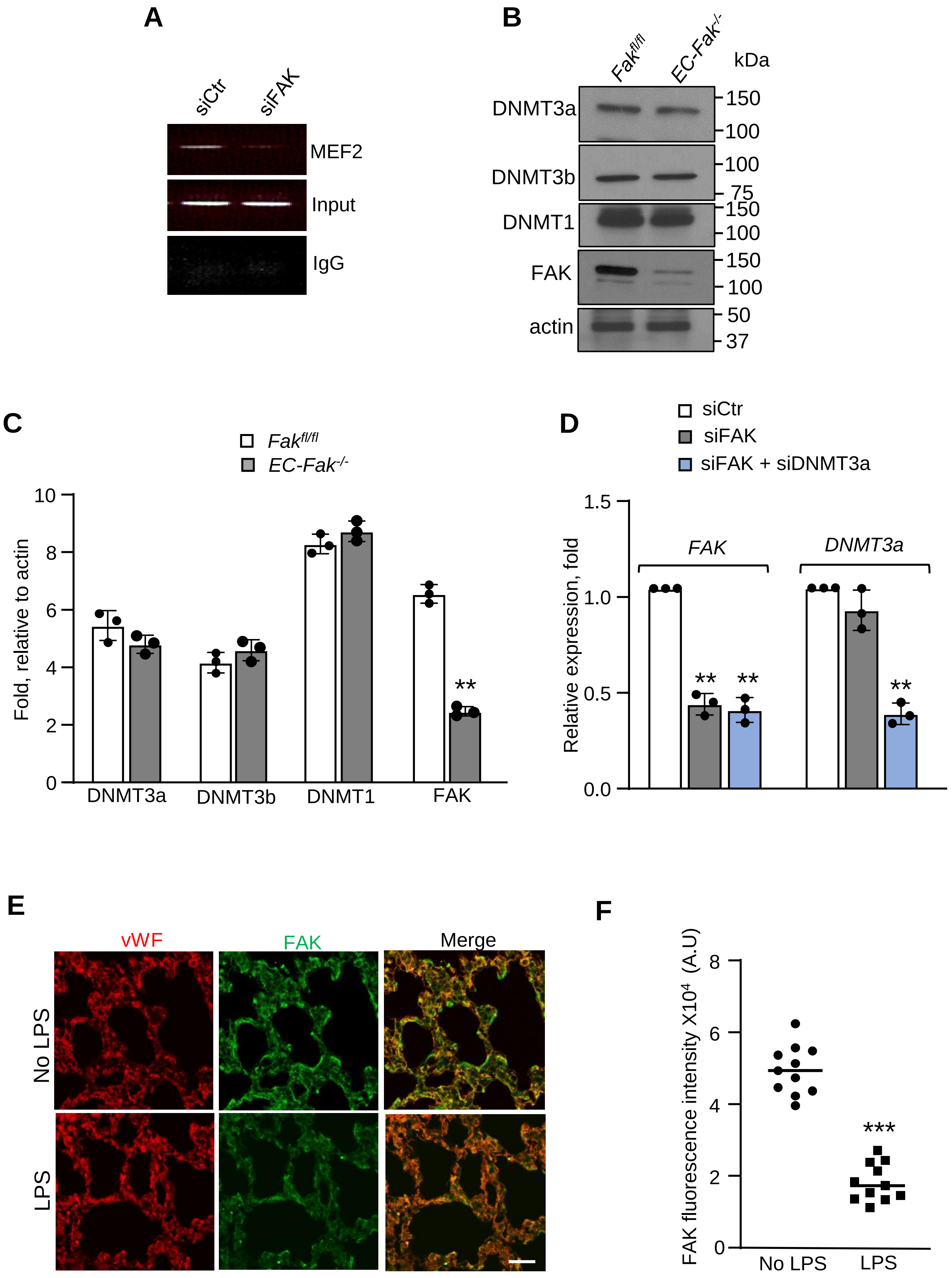

#### Supplementary 4

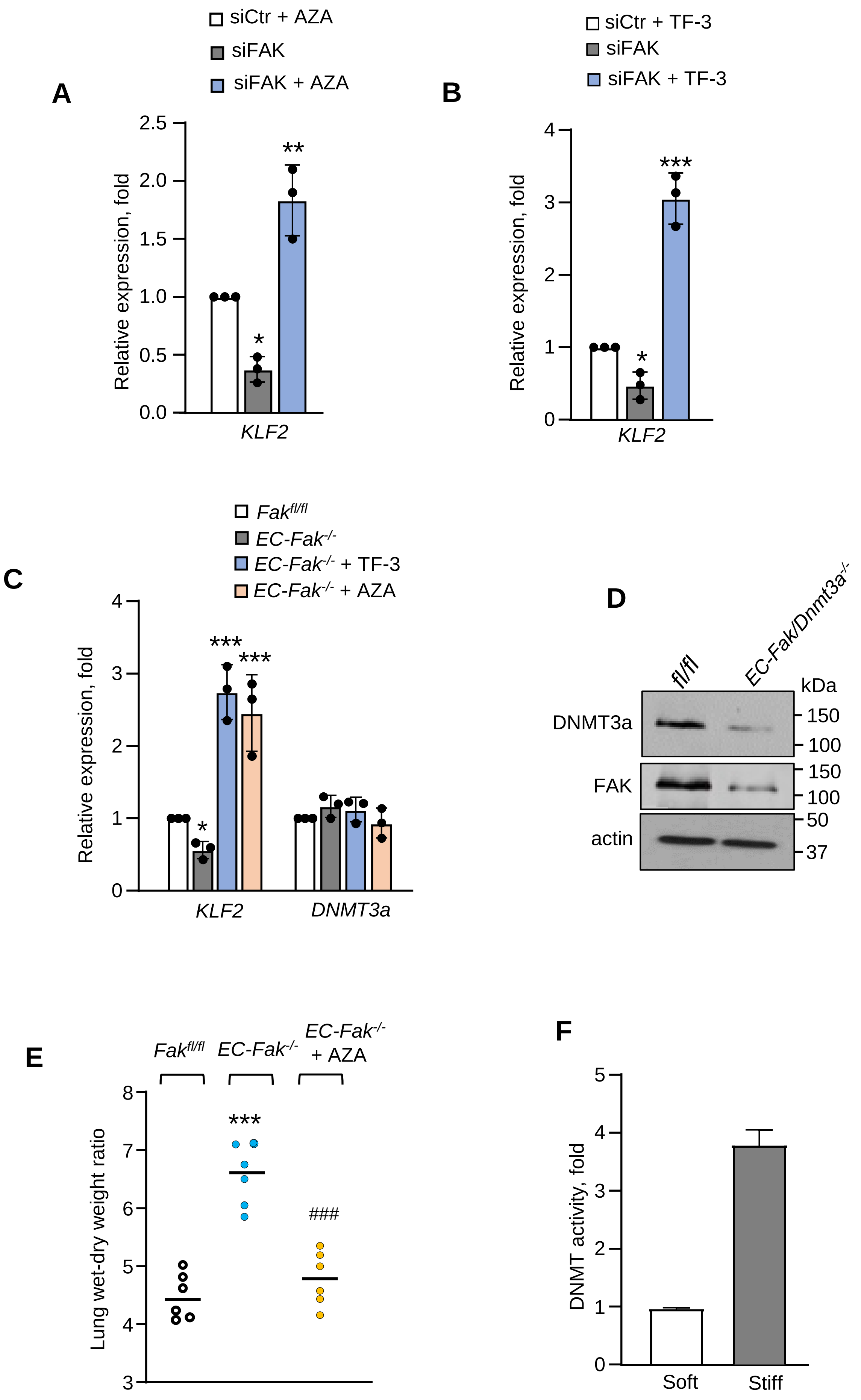

#### Supplementary 5

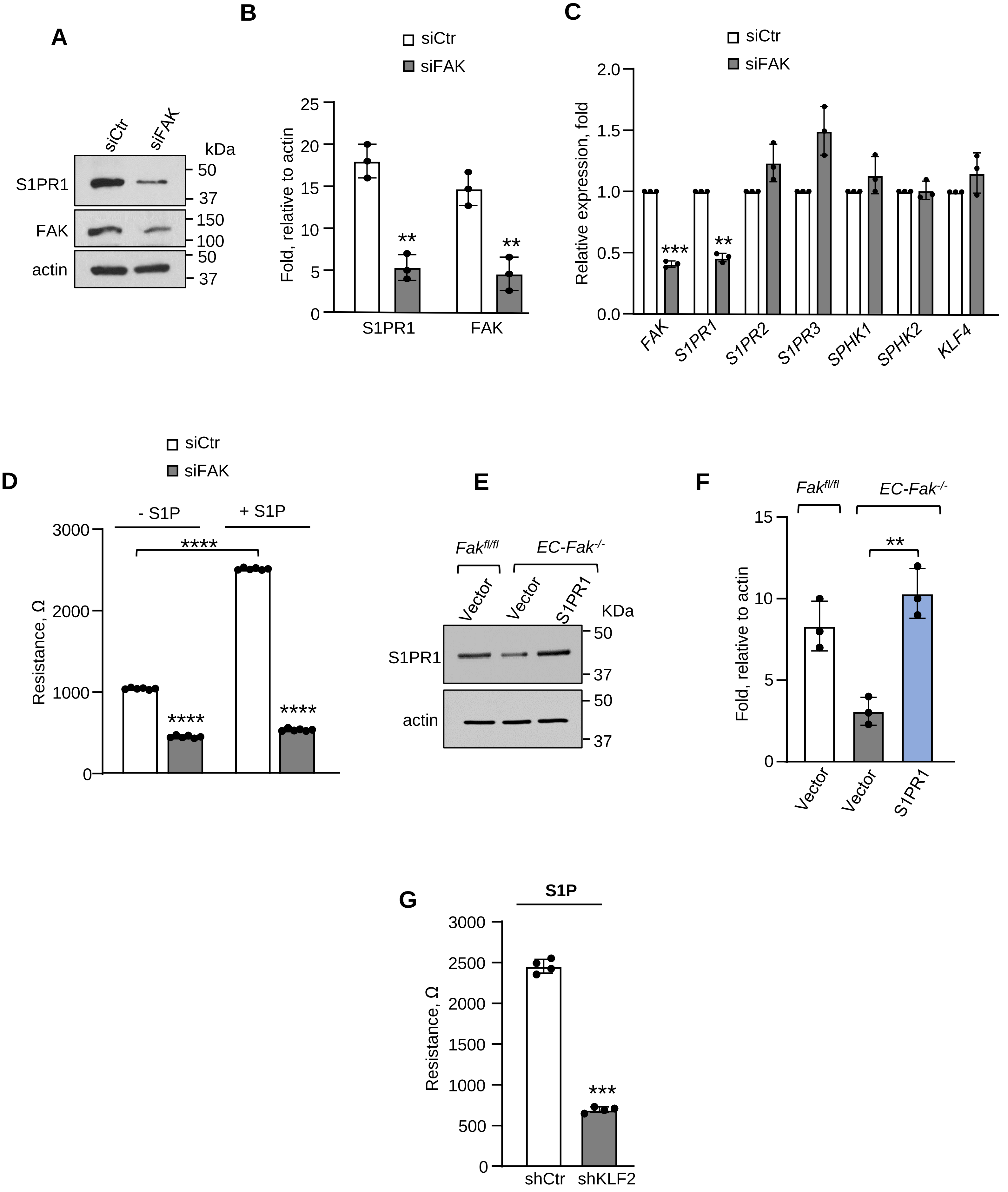

**Supplementary 6**

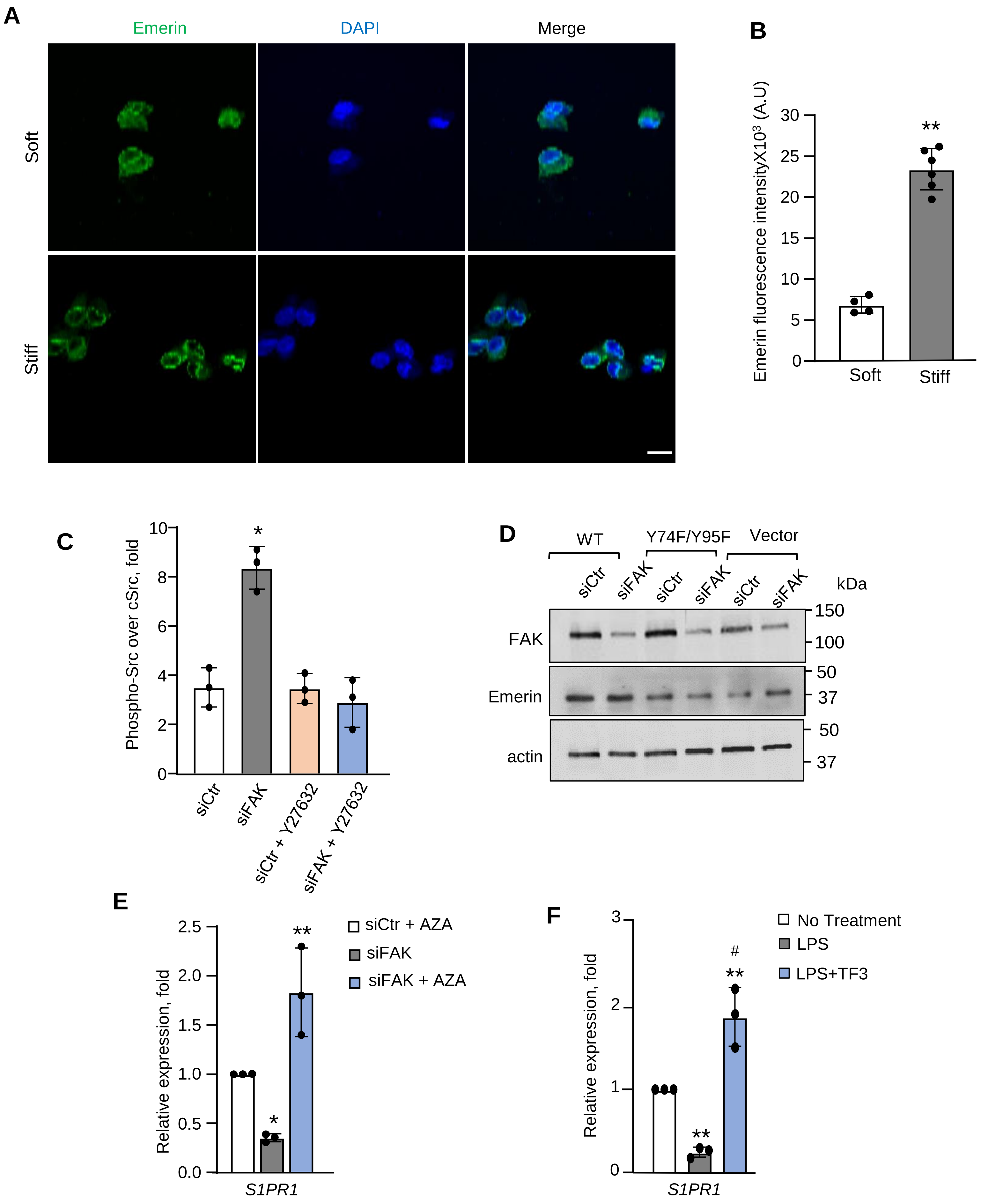
